# Cingulate-motor circuits update rule representations for sequential choice decisions

**DOI:** 10.1101/2021.05.26.445888

**Authors:** Daigo Takeuchi, Dheeraj Roy, Shruti Muralidhar, Takashi Kawai, Chanel Lovett, Heather A. Sullivan, Ian R. Wickersham, Susumu Tonegawa

## Abstract

Anterior cingulate cortex mediates the flexible updating of an animal’s choice responses upon rule changes in the environment. However, how anterior cingulate cortex entrains motor cortex to reorganize rule representations and generate required motor outputs remains unclear. Here, we demonstrate that chemogenetic silencing of the projection terminals of cingulate cortical neurons in secondary motor cortex disrupted sequential choice performance in trials immediately following rule switches, suggesting that these inputs are necessary to update rule representations for choice decisions stored in the motor cortex. Indeed, the silencing of cingulate cortex decreased rule selectivity of secondary motor cortical neurons. Furthermore, optogenetic silencing of cingulate cortical neurons that was temporally targeted to error trials immediately after rule switches exacerbated errors in following trials. These results suggest that cingulate cortex monitors behavioral errors and update rule representations in motor cortex, revealing a critical role for cingulate-motor circuits in adaptive choice behaviors.

## Main

A central feature of animal intelligence is the hierarchical organization of behaviors and the capacity to adopt flexible strategies that allow complex sequential behaviors^1–3^. Previous studies in primates suggested that premotor cortex and supplementary motor areas underlie planning and execution of sequential movements^4–7^. More recently, it has been demonstrated that neurons in the rodent secondary motor cortex (M2) similarly code initiation of sequential movements, motor planning, memory for upcoming choice actions and values for choice actions, suggesting that they maintain representations of sensorimotor associations for adaptive choice behaviors^8–14^. These findings raise a question of what circuit mechanisms enable M2 neurons to process context-dependent information such as task rules. More specifically, how is this information provided from input brain regions and how are they processed by M2 circuits?

Studies using humans and animals have shown that medial prefrontal areas including anterior cingulate cortex (ACC) mediates flexible decisions in the face of rule changes in the environment (e.g., task switching) or under uncertain conditions^15–20^. However, despite abundant anatomical evidence of cingulate projections to motor cortices^21–24^, how anterior cingulate cortex entrains motor cortex to update neural representations for rules upon sudden rule changes in the environment remains poorly understood. In this study, we addressed this issue by examining what aspects of choice behaviors are mediated by the ACC→M2 circuit and how ACC circuits modulate neural activity in M2 after rule switches in which mapping of choice actions and rewards are suddenly changed. Our results suggest that anterior cingulate circuits monitor behavioral errors and reorganize choice behaviors upon sudden rule switches by updating rule representations in motor cortex.

## Results

### Conditional action sequencing task

We devised a novel behavioral task in which an animal updates its sequential choice response for rewards upon abrupt rule changes (hereafter referred to as the “conditional action sequencing task” or CAS task). Briefly, in the CAS task, animals were required to choose left or right ports based on an auditory tone cue stimulus. Under the “1 step rule” condition, animals received a reward after correctly poking a left or right port instructed by one of two tone cues, and could start the next trial after an inter trial interval (ITI) (Fig. 1, top schematic). When animals poked an incorrect side port, it received no reward and instead a buzzer tone was delivered. Under the “2 steps rule” condition, animals received a first reward after making a correct first choice and then received another reward after poking the opposite side port (Fig. 1, bottom schematic). If the animal pushed the center lever before choosing the opposite side port, they received no reward and instead heard the buzzer tone. Rules were switched between these two conditions every 55 trials, requiring the animals to adjust their choice behavior driven by water rewards or buzzer penalties (Extended Data Fig. 1). We trained rats to achieve over 70% success rate for both 1 step and 2 steps rule conditions, and for both of the tone cues.

**Fig. 1.**
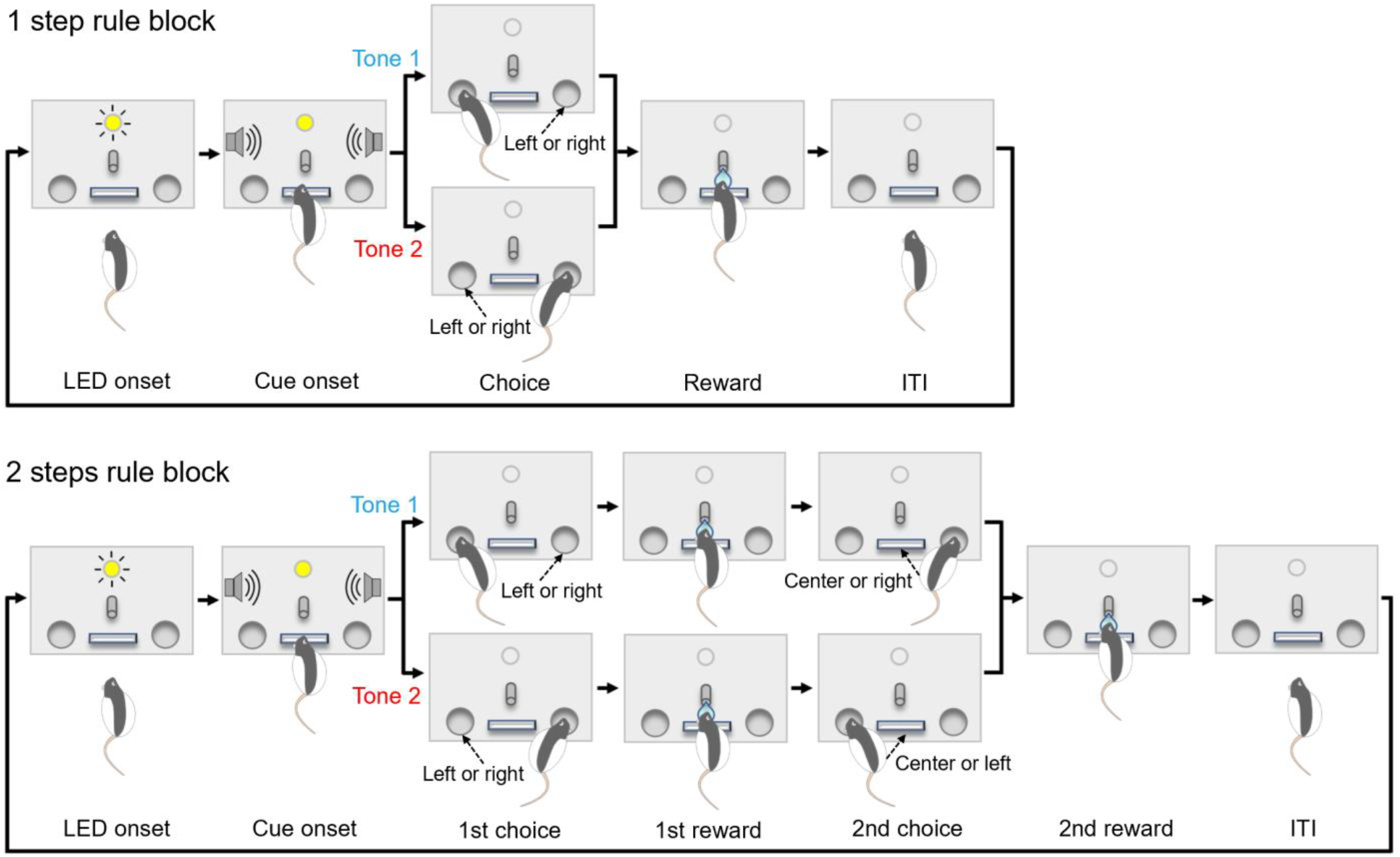
Conditional action sequencing task. Rats were trained and tested in a chamber in which a lever, a water spout and an LED were installed on front wall with two infrared (IR) ports being on left and right sides. Sound speakers were equipped on side walls. Task rules were switched between 1 step rule and 2 steps rule conditions every 55 trials. LED onset signals the end of inter trial interval (ITI) and rats can start a new trial by pushing the center lever. In 1 step rule condition (top), animals received a water reward after correctly poking left or right IR port instructed by one of two tone cues. When animals poked an incorrect port, it received no reward and instead a buzzer sound was delivered with an extra waiting being imposed in ITI before starting the next trial (“Error” trial). In 2 steps rule condition (bottom), animals received a reward after making a correct first choice and then received another reward after poking the opposite side port. If animal made an incorrect first choice, no reward was delivered. Instead a buzzer sound was delivered and an extra waiting time was imposed in ITI before starting the next trial (“Error 1” trial). If animal made a correct first choice but pushed the center lever before poking the opposite side port, it received no reward. Instead a buzzer sound was delivered and an extra waiting time was imposed in ITI (“Error 2” trial).

### Silencing anterior cingulate excitatory neurons disrupted the animal’s ability to adapt to rule switches

To examine the potential roles of ACC in animal’s flexibly adapting to rule switches in the CAS task, we chemogenetically silenced neural activity in ACC and examined its effect on animal’s task performance. First, we bilaterally injected an inhibitory DREADD virus in the ACC of rats that had been trained on the CAS task (Fig. 2a and Extended Data Fig. 2a). Intraperitoneal administration of clozapine-N-oxide (CNO) decreased spiking activity in ACC, an effect that lasted over 60 minutes after reaching the plateau level of firing (Extended Data Fig. 2b). After animals recovered from virus injection surgeries, we injected CNO or saline and tested their 2 steps choice performance for one block followed by another block after a rule switch. To quantify the animal’s ability to adapt to rule switches from 1 step to 2 steps rules, we measured the animal’s second choice performance (% Error 2) (see Extended Data Fig. 1d for details). Similarly, to quantify the animal’s ability to adapt to rule switches from 2 steps to 1 step rules, we measured frequency of non-rewarded 2nd action (Nr2A) (Extended Data Fig. 1d). In a representative saline session (Fig. 2b,c and Extended Data Figs. 3 and 4), second choice errors (Error 2) were observed only in the first few trials after rule switches, indicating that the animal could adapt to rule switches from 1 step to 2 steps rules within a few trials (Fig. 2b, thick blue line. Also see right panels in Extended Data Fig. 3a,b). In contrast, when the same animal received CNO in another session, second choice errors persisted beyond trials that immediately followed rule switches from 1 step to 2 steps rules (Fig. 2b, thick pink line. Also see left panels in Extended Data Fig. 3a,b), suggesting that silencing ACC neurons impaired the animal’s ability to adapt to rule switches from 1 step to 2 steps rules. Similarly, the animal showed Nr2A more frequently in trials that immediately followed rule switches from 2 steps to 1 step, which decreased within 10-20 trials after rule switches (Fig. 2c, blue), indicating that the animal could adapt to rule switches from 2 steps to 1 step conditions (Fig. 2c and Extended Data Fig. 3a). Same tendency was observed in CNO session (Fig. 2c), suggesting that silencing ACC neurons did not impact the animal’s ability to adapt to rule switches from 2 steps to 1 step rules. Response times for 1st and 2nd choices in the CNO condition were greater than those in the saline condition (Extended Data Fig. 4e,f). We repeated these experiments using eleven rats injected with the inhibitory DREADD virus in ACC. Results showed that the 2nd choice error rate (%Error 2) in 4 the 2 steps condition was significantly greater for the CNO condition (Fig. 2d. Also see Extended Data Fig. 5a for individual animals’ data). In contrast, no difference was observed in the 1st choice performance in the 2 steps condition or in that of the 1 step condition (Extended Data Fig. 5b,c), indicating that chemogenetic silencing of ACC excitatory neurons did not affect the animals’ ability to discriminate between the two auditory cues or their ability to make choice responses. To examine if chemogenetic silencing of ACC neurons impaired the animals’ ability to make sequential actions or the ability to adjust their choice behavior according to rule changes, we calculated the 2nd choice performance (%Error 2) in 2 steps rule for the 1st block of the session (i.e., non-rule switching block) and for subsequent Rule Switch-blocks separately. We found a significant difference in the error rate between CNO and saline conditions for Rule Switch blocks while there was no significant difference for the 1st block (Fig. 2e,f). This suggests that the silencing of ACC neural activity affected the animals’ ability to adjust its choice responses to rule changes from 1 step to 2 steps more severely than its ability to make sequential actions. We next split the 2 steps block into three epochs and compared 2nd choice performance among these epochs. We found that, in the saline condition, animals committed 2nd choice errors more frequently in 1st epoch (i.e., trials that immediately followed rule switches) than in 2nd or 3rd epochs (Fig. 2h, blue). On the other hand, the higher error rate persisted beyond 1st epoch in the CNO condition (Fig. 2h, red). This effect was observed in Rule Switch blocks (Fig. 2h) but not in the 1st block (Fig. 2g), suggesting that silencing of ACC neural activity impaired the animal’s ability to adjust choice responses upon rule switches from 1 step to 2 steps (also see Extended Data Fig. 5f,g for individual animals’ data). We further divided the 1st epoch (i.e., the first 18 trials immediately after rule switches) into three periods (1-6th, 6-12th, 13-18th trials) and compared the animals’ 2nd choice performance between CNO and saline conditions (Fig. 2i). We found a significant difference in the 2nd choice performance between CNO and saline conditions in the 3rd period of the 1st epoch (i.e., corresponding to the 13-18th trials after rule switches), indicating that, on average, the impairment in adjusting choice responses to rule changes from 1 step to 2 steps due to the silencing of ACC neural activity showed up within the first 10-20 trials in the 2 steps rule block. mCherry control animals did not show any significant difference in the 2nd choice performance between CNO and saline conditions, excluding the possibility that the observed effect of chemogenetic silencing on task performance in previous experiments was caused by CNO administration itself (Extended Data Fig. 6).

**Fig. 2.**
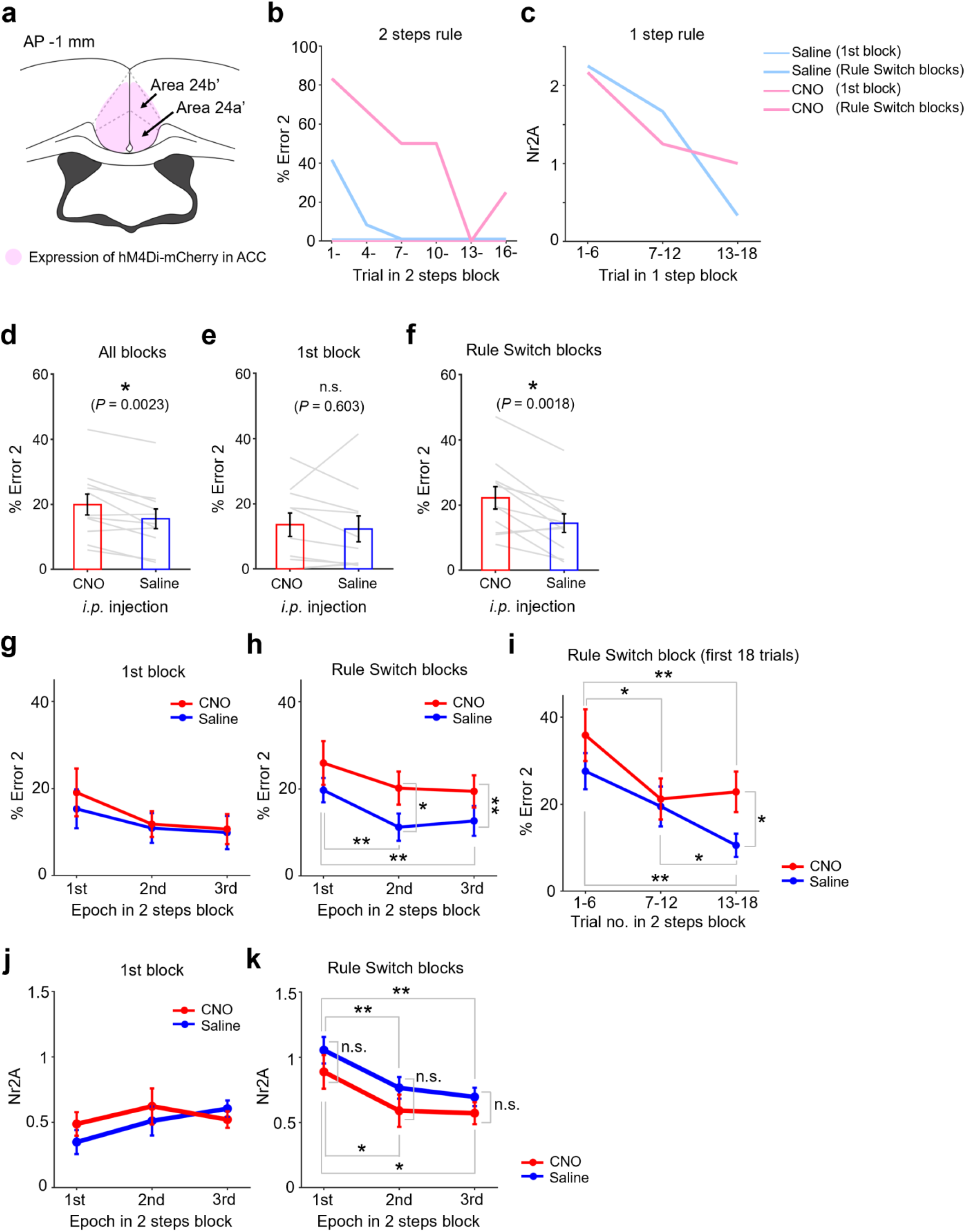
Chemogenetic silencing of ACC neurons disrupted the animals’ sequential choice performance after rule switches. **a,** Rats were injected with AAV5-CaMKIIa-hM4Di-mCherry virus in ACC (area 24a’/24b’). At least three weeks after virus injection, animals were tested on CAS task with *i.p.* injections of either saline or clozapine-N-oxide (CNO) solutions. Black filled structure is lateral ventricle. **b**, CAS task performance in two representative sessions with an *i.p.* injection of CNO solution (10 mg/kg) or of saline solution. 2nd choice performance in 2 steps rule condition (%Error 2) were plotted against trial no. in the block (see Extended Data Figs. 3 and 4 for further details of task performance in these representative sessions). The first 18 trials in 2 steps rule blocks were grouped into 6 periods consisting of 3 trials and the average task performance for each period was plotted. Pink, CNO solution. Blue, saline. Thin lines, 1st block. Thick lines, Rule Switch blocks. **c**, Non-reward 2nd action (Nr2A) in 1 step rule condition for the same two representative sessions were plotted against trial no. in the block. The first 18 trials in 1 step rule block were grouped into 3 periods consisting of 6 trials and the average task performance for each period was plotted. Note that there was no “1st block” in 1 step rule condition because the sessions started with 2 steps rule condition, thus no thin blue/pink line in **c** (also see Extended Data Figs. 3 and 4). **d**, Group result of 2nd choice performance in 2 steps rule condition (%Error 2) with *i.p.* injections of saline or CNO solutions. For CNO data, sessions with doses of 10 mg/kg and 20 mg/kg were combined (see Extended Data Fig. 6a for individual animals’ data with each CNO dose shown separately). Performance for two tone cues were averaged. Paired *t*-test, n = 11 rats. Error bars, s.e.m. **e,** Same format as in **d**, but % Error 2 for trials in 1st block (i.e., non-rule switching block). **f,** Same format as in **d** and **e**, but % Error 2 for trials in Rule Switch blocks. **g,** 2nd choice performance (%Error 2) was plotted separately for three epochs in 1st block. Blue, saline. Red, CNO. For CNO data, sessions with doses of 10 mg/kg and 20 mg/kg were combined (see Extended Data Fig. 6f,g for results with each CNO dose shown separately). Performance for two tone cues were averaged. Repeated measures two-way analysis of variance (ANOVA) with both CNO dose and epoch being within-subject factors revealed no main effect of CNO dose or epoch (*P* > 0.5 for CNO dose and *P* > 0.2 for epoch, n = 11 rats). Error bars, s.e.m. **h,** Same as in **g**, but for Rule Switch blocks. CNO dose but not epoch showed a significant main effect (*P* = 0.017, F_1,62_ = 6.02 for dose; *P* = 0.098, F_2,62_ = 2.41 for epoch). No interaction was detected (*P* > 0.8). Post-hoc comparisons were conducted using paired *t*- test with Bonferroni’s correction across epochs. **, *P* < 0.01. *, *P* < 0.05. **i,** 2nd choice performance in 2 steps rule condition (%Error 2) was plotted separately for 1-6th, 7-12th and 13-18th trials of Rule Switch blocks. **, *P* < 0.01. *, *P* < 0.05. Paired *t*-test, n = 11 rats. Error bars, s.e.m. **j,** Average number of non-rewarded 2nd action (Nr2A) per trial was plotted separately for three epochs in Rule Switch blocks of 1 step rule condition. Repeated measures two-way ANOVA with both CNO dose and epoch being within-subject factors revealed no main effect of CNO dose or epoch (*P* > 0.6 for CNO dose and *P* > 0.4 for epoch). **k,** Same as in **j**, but for Rule Switch blocks. Significant main effect was detected for epoch (*P* = 0.0022, F_2,62_ = 6.80) but not for dose (*P* = 0.061). Post-hoc comparisons were conducted using paired *t*-test with Bonferroni’s correction across epochs. **, *P* < 0.01. *, *P* < 0.05. n = 11 rats. Error bars, s.e.m.

We next compared the average number of non-rewarded 2nd action (Nr2A) per trial across all three epochs in the 1 step rule condition. Results showed a significant effect of epoch in Rule Switch blocks but not in the 1st block (Fig. 2j,k), indicating that animals could adapt to rule switches from 2 steps to 1 step rules (also see Extended Data Fig. 5d,e). Interestingly, no effect was found for CNO dose, suggesting that, while the silencing of neural activity in ACC impaired animals’ ability to adapt to rule switches from 1 step to 2 steps rules, it did not affect the ability to adapt to rule switches in the opposite direction (i.e., from 2 steps to 1 step rules) (also see Extended Data Fig. 5h,i for individual animals’ data).

### Silencing ACC neuronal terminals in M2 disrupted the animals’ ability to adapt to rule switches

Given the abundant anatomical evidence of projections from ACC to motor cortices^21–24^, we hypothesized that projections from ACC to motor cortices entrain motor cortex to update neural representations for rules that are maintained in motor circuit for generating required motor outputs. We first tried to identify anatomical projections from ACC to motor cortices using a genetically modified rabies virus approach (Fig. 3a)^25^. We found that a posterior portion of M2 (an area referred to as “frontal orienting field (FOF)” in previous studies and hereafter referred to as “M2”)^8, 26^ received projections from ACC (area 24a’/24b’) (Fig. 3b,c and Extended Data Fig. 7). We also validated these projections from ACC to M2 using AAVretro-Cre and AAV-DIO viruses (Fig. 3d,e).

**Fig. 3.**
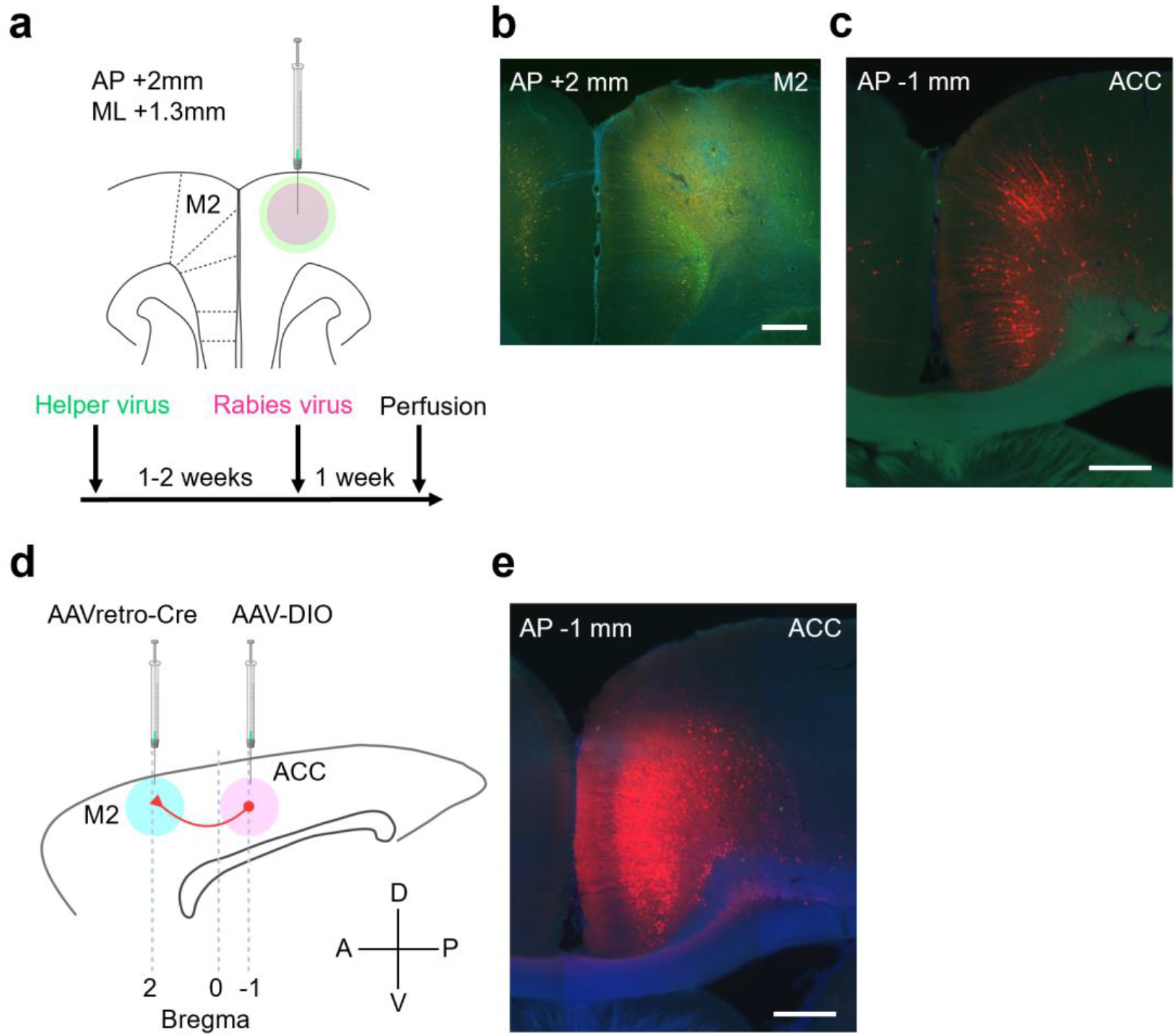
Anatomical projections from ACC to M2. **a,** Anatomical projections from ACC to M2 were visualized using a genetically modified rabies virus system. 1-2 weeks after helper virus injection in M2 (a cocktail solution of AAV1-synP-FLEX-sTpEpB and pENN.AAV.CaMKII.0.4.Cre.SV40), rabies virus (RVΔG-4mCherry) was injected at the same coordinate. **b,** A coronal section of the virus injection site in M2 (this is a magnified view of a region pointed by red arrow in panel no. 4 in Extended Data Fig. 7a). Neurons infected by helper virus expressed GFP. Scale bar, 0.5 mm. **c,** ACC neurons that were retrogradely infected with rabies virus expressed mCherry. Scale bar, 0.5 mm. **d,** Projections from ACC to M2 were validated using AAVretro virus. Sagittal view of rat brain. Rats were injected with AAVretro-pmSyn1-EBFP-cre and AAV5-hSyn-DIO-hM4Di-mCherry viruses in M2 and ACC, respectively. **e,** ACC neurons that were infected with AAVretro virus expressed mCherry after Cre recombination. Scale bar, 0.5 mm.

We then asked what aspects of choice behaviors are mediated by the ACC→M2 circuit that we identified. More specifically, we wanted to examine the potential roles of projections from ACC to M2 in the animal’s ability to update sequential choices upon rule switches. Using bilateral infusion cannulae targeting M2 in rats that had been injected with the inhibitory DREADD virus in ACC, we infused CNO solution in M2 and examined its effect on task performance (Extended Data Fig. 8a,b). The 2nd choice error rate in the 2 steps condition showed an increased trend in the CNO condition as compared to saline control (Fig. 4a). In contrast, no difference was observed in the 1st choice performance in the 2 steps condition or in that of the 1 step condition (Extended Data Fig. 8c,d). We separately calculated 2nd choice performance in the 2 steps condition for the 1st block (i.e., non-rule switching block) and for subsequent Rule Switch-blocks. The 2nd choice error rate was greater in the CNO condition relative to saline in Rule Switch blocks but not for the 1st block (Fig. 4b,c), suggesting that chemogenetic silencing of ACC terminals in M2 impaired the animals’ ability to adjust its choice responses to rule changes from 1 step to 2 steps more severely than its ability to make sequential actions. We next split the 2 steps rule blocks (55 trials) into three epochs (18, 18 and 19 trials for 1st, 2nd and 3rd epochs, respectively) and compared the 2nd choice performance across epochs. Results showed a significant effect of CNO in Rule Switch blocks while no such effect was found in the 1st block (Fig. 4d,e). We further divided the 1st epoch (i.e., the first 18 trials immediately after rule switches) in Rule Switch blocks into three periods (6 trials each) and compared the animals’ 2nd choice performance between CNO and saline conditions (Fig. 4f). We found a significant difference in the 2nd choice performance between CNO and saline conditions in the 3rd period (i.e., corresponding to the 13-18th trials after rule switches), indicating that, similar to the *i.p.* injection experiments (Fig. 2i), the impairment in adjusting choice responses to rule changes showed up within the first 10-20 trials in the 2 steps rule block. We also compared the frequency of non-rewarded 2nd actions across all three epochs in the 1 step-block. Similar to the results obtained in *i.p.* injection experiments (Fig. 2k), a decreasing tendency of Nr2A across epochs in Rule Switch blocks of the 1 step rule condition was observed but no difference was found in the Nr2A frequency between CNO and saline conditions in either of the three epochs (Fig. 4g). These results suggested that the silencing of ACC terminals in M2 impaired the animals’ ability to adjust their choice responses upon rule switches from 1 step to 2 steps rules but did not affect their ability to adapt to rule switches in the opposite direction (i.e., from 2 steps to 1 step rules) (also see Extended Data Fig. 8e). Finally, we conducted chemogenetic silencing of prelimbic/infralimbic cortex and ventral thalamic nuclei. We found no effect in any aspect of the task performance in both experiments (Extended Data Figs. 9 and 10 for results of prelimbic/infralimbic cortex and of ventral thalamic nuclei, respectively), indicating that ACC and its projections to M2, but not prefrontal cortex or ventral thalamic nuclei are specifically recruited for reorganizing sequential choice decisions upon rule switches.

**Fig. 4.**
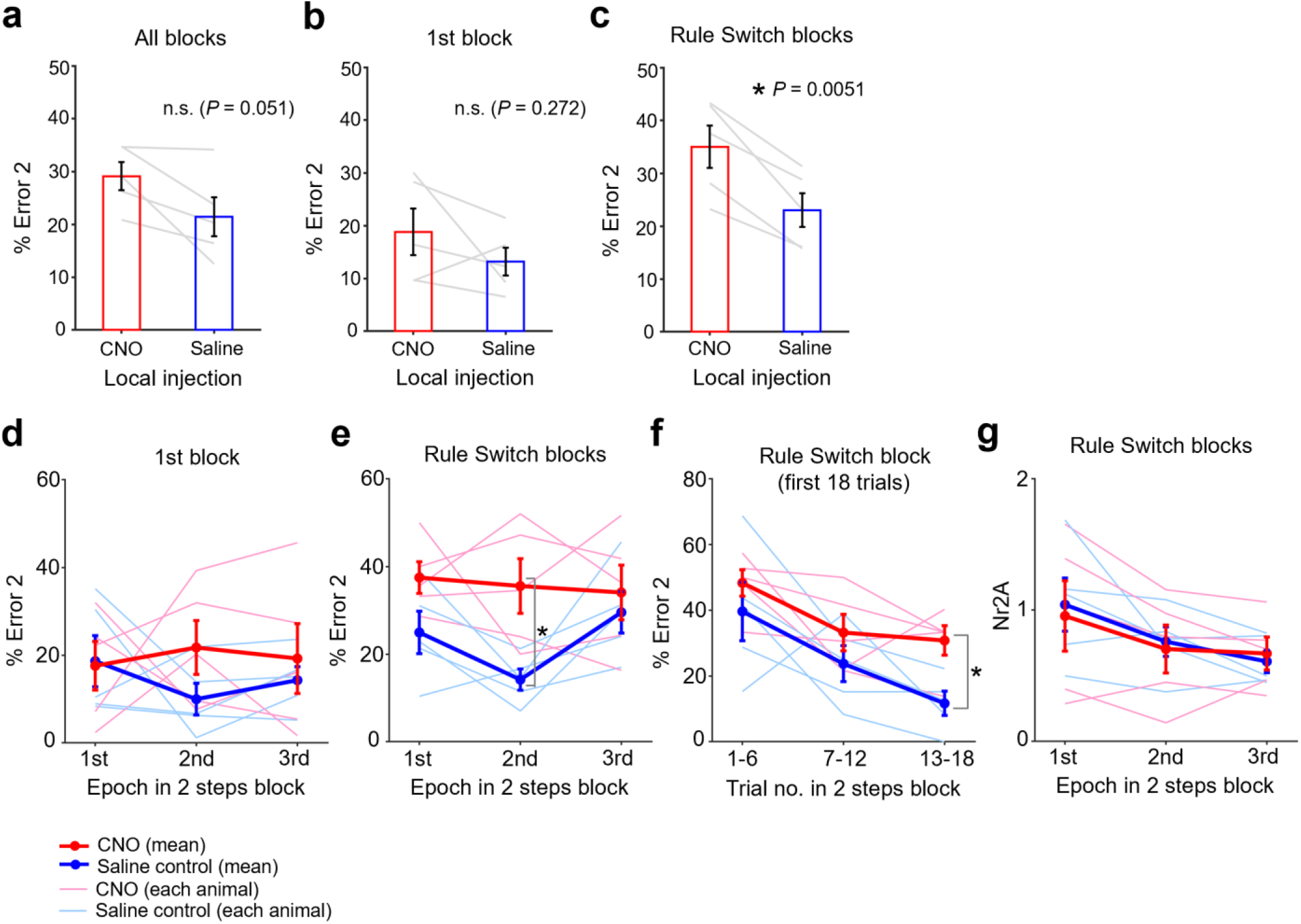
Chemogenetic silencing of ACC neuronal terminals in M2 disrupted the animals’ sequential choice performance after rule switches. **a,** Group result of 2nd choice performance in 2 steps rule condition (%Error 2) with a local infusion of either saline or CNO solution. **b**, Same as in **a**, but % Error 2 for trials in 1st block (i.e., non-rule switching block). **c**, Same as in **a** and **b**, but % Error 2 for trials in Rule Switch blocks. **d**, 2nd choice performance (%Error 2) was plotted separately for three epochs (“1st”, “2nd” and “3rd”) in 1st block for sessions with a local infusion of saline or CNO solution. Thick red and blue lines represent across-animal averages of CNO and saline conditions, respectively (n = 5 rats). Thin lines represent individual animals. 1st, 2nd and 3rd epochs correspond to 1-18th, 19-36th and 37-55th trials. Neither CNO dose nor epoch in 2 steps rule block showed no main effect (*P* > 0.2 for CNO dose and *P* > 0.9 for epoch). **e**, Same format as in **d**, but for Rule Switch blocks. CNO dose showed a significant main effect (*P* = 0.00402, F_1,26_ = 9,969) while epoch did not (*P* > 0.3). Post-hoc comparisons were conducted using paired *t*-test with Bonferroni’s correction across epochs. *, *P* = 0.0483. **f**, 2nd choice performance in 2 steps rule condition (%Error 2) was plotted separately for 1-6th, 7-12th and 13-18th trials in 1st epoch of Rule Switch blocks. *, *P* = 0.0432. Paired *t*-test, n = 5 rats. **g**, Average number of non-rewarded 2nd action (Nr2A) per trial was plotted separately for three epochs in Rule Switch blocks of 1 step rule condition. Repeated measures two-way ANOVA with both CNO dose and epoch being within-subject factors revealed no main effect of CNO (*P* > 0.8). Epoch showed a moderate effect but did not reach statistical significance (*P* = 0.105, F_2,26_ = 2.461). n = 5 rats. Error bars, s.e.m.

### Chemogenetic suppression of ACC neural activity decreased rule selectivity in M2 neuron

We next asked how ACC circuits affect neural activity in M2. We unilaterally implanted array electrodes in M2 and measured spiking activity while the animals were performing the CAS task (CNO or saline control conditions) (Fig. 5a). We obtained 900 single-units in 43 sessions from five rats (594 and 306 units in saline and CNO conditions, respectively). Without CNO, some neurons showed activity that was selective to a specific rule (i.e., 1 step or 2 steps rule) even before the animal made 1st choice responses (Fig. 5b,c and Extended Data Fig. 11a,b. Also see Fig. 5d,e for an example single-unit activity with an *i.p.* injection of CNO). We calculated mean firing rate of M2 single-units during a 1 sec period immediately before animals made their 1st choice (“pre-choice period”) and found that chemogenetic suppression of ACC neural activity decreased firing rate both in ipsilateral and contralateral choice conditions (Fig. 5f).

**Fig. 5.**
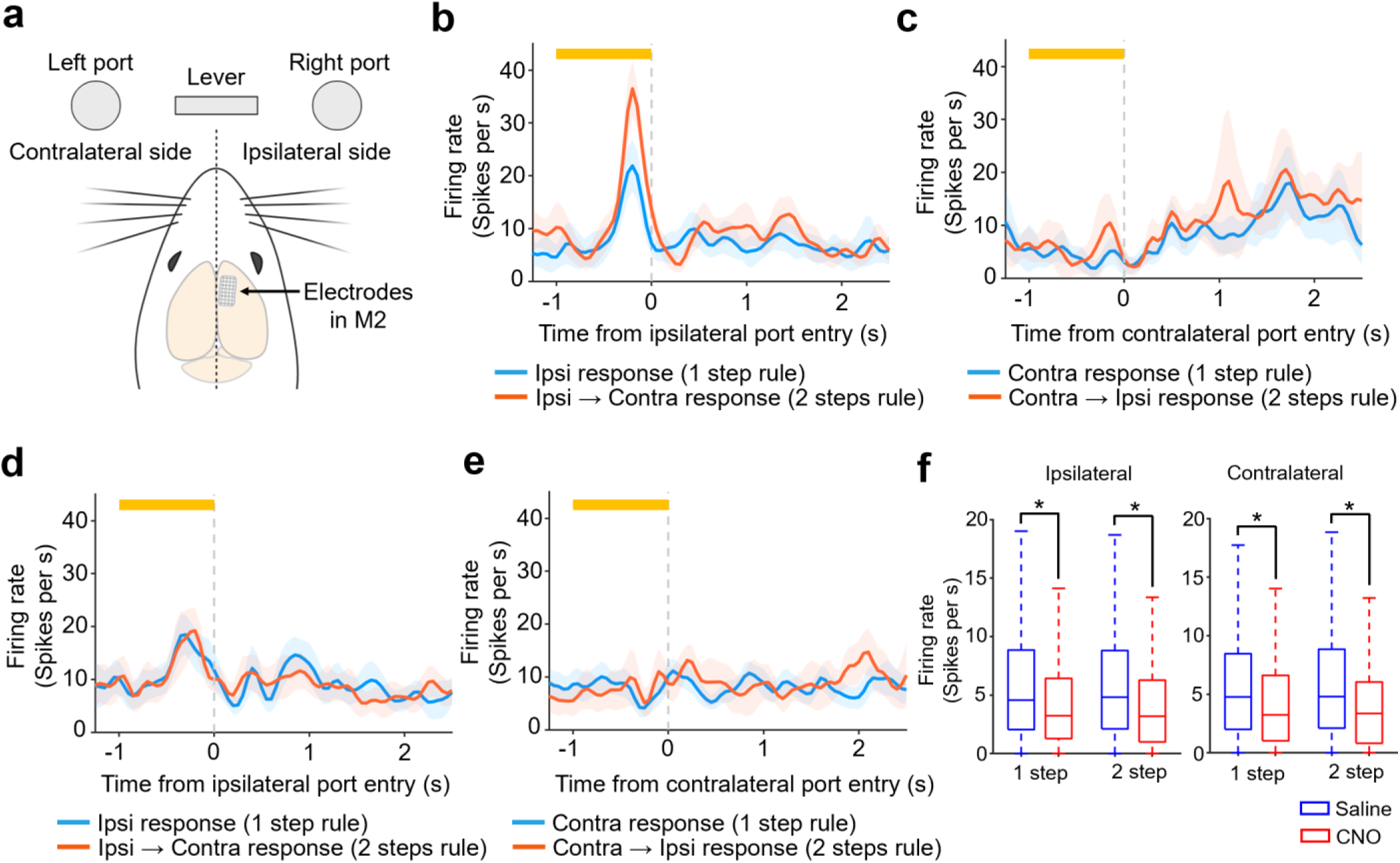
Chemogenetic silencing of ACC decreased firing rate in M2 neurons. **a,** Ipsilateral and contralateral sides viewed from a rat’s cerebral hemisphere in which M2 single-unit activity was measured. Rats were implanted with Utah array electrodes in M2 of either left or right hemisphere and neural activity was measured during their task performance with *i.p.* injections of saline or CNO solutions (20mg/kg). In the following analysis, trials were classified according to which side port (ipsilateral or contralateral) rats chose as their choices (1 step rule condition) or their 1st choices (2 steps rule condition). **b,** Peri-event time histogram (PETH) of a representative single-unit showing rule selective responses before making a correct response to the side port that was located on the ipsilateral side of neural activity measurements (“ipsilateral condition”). In 1 step rule block (blue line), the rat made a choice to the ipsilateral side port while, in 2 steps rule block (red line), the rat made a 1st choice to the ipsilateral side port and then made a 2nd choice to the contralateral side port. Neural activity measured in trials of Rule Switch blocks were plotted (note that neural activity in 1st block of the session was not included). Orange bar at the top, a 1 sec period immediately before rat’s entry to the ipsilateral side port (pre-choice period). Shaded bands, 95% confidence intervals. **c,** Same as in b, but PETH was calculated using trials in which the rat made a correct response to the side port that was located on the contralateral side of neural activity measurements (“contralateral condition”). **d**, PETH of a single-unit measured in a session in which the rat received an *i.p.* injection of CNO solution. **e**, Same as in **d**, but for contralateral condition. **f,** Population result of firing rate during the pre-choice period of 1 step and 2 steps rule conditions. Box-and-whisker plots indicate the minimum, 25th, 50th, 75th percentiles, and maximum excluding outliers (i.e., 1.5 times greater than the interquartile range). *, *P* < 10^-5^; Mann-Whitney (n = 594 and n = 306 single-units for saline and CNO conditions, respectively).

We then quantified rule selectivity during the pre-choice period for each single-unit using receiver operating characteristic (ROC) analysis^27^, which measures the degree of overlap between two response distributions^28, 29^. For each M2 single-unit the preferred and non-preferred rule conditions were compared, given two distributions of neuronal activity (see Methods). An ROC curve was then generated by taking the observed firing rate of a neuron and then the area under the ROC curve was calculated. A value of 0.5 indicates that the two distributions were completely overlapped, and thus the neuron is not selective to the rules. A value of 1.0, on the other hand, indicates that the two distributions were completely separated and so the neuron is very selective. Time course of rule selectivity of the representative M2 single-unit showed an increase of rule selectivity in the pre-choice period in ipsilateral choice condition but not in contralateral condition (Fig. 6a,b. Also see Extended Data Fig. 11c,d). We repeated the same analysis for all the M2 single-units that exhibited mean firing rate greater than 3 Hz during pre-choice period in either ipsilateral or contralateral conditions (437 and 195 single-units for saline and CNO conditions, respectively) (see Methods). Population-averaged time course of rule selectivity showed that chemogenetic silencing of ACC decreased rule selectivity of M2 neurons in “ipsilateral” trials and this tendency could be seen not only after animals made ipsilateral choice but even before making the choice (Fig. 6c). Such an effect was not seen in contralateral conditions (Fig. 6d). We split the Rule Switch blocks into three epochs and examined rule selectivity in each epoch for both CNO and saline conditions. The administration of CNO decreased rule selectivity in trials in which animals made a 1st response to the ipsilateral side followed by a 2nd response to the contralateral side (Fig. 6e, top left panel), while such a tendency was not observed in trials in which animals made its 1st response to the contralateral side followed by the 2nd choice to the ipsilateral side (Fig. 6e, top right and bottom right panels). The effect observed in the ipsilateral condition was greatest in the 1st epoch that immediately followed rule switches from the 1 step to 2 steps rules (Fig. 6e, top left panel). These results suggest that, immediately after rule switches from 1 step to 2 steps conditions, ACC modulates neural activity in M2 that encodes rule representations for specific sequential choices starting from ipsilateral to contralateral sides.

**Fig. 6.**
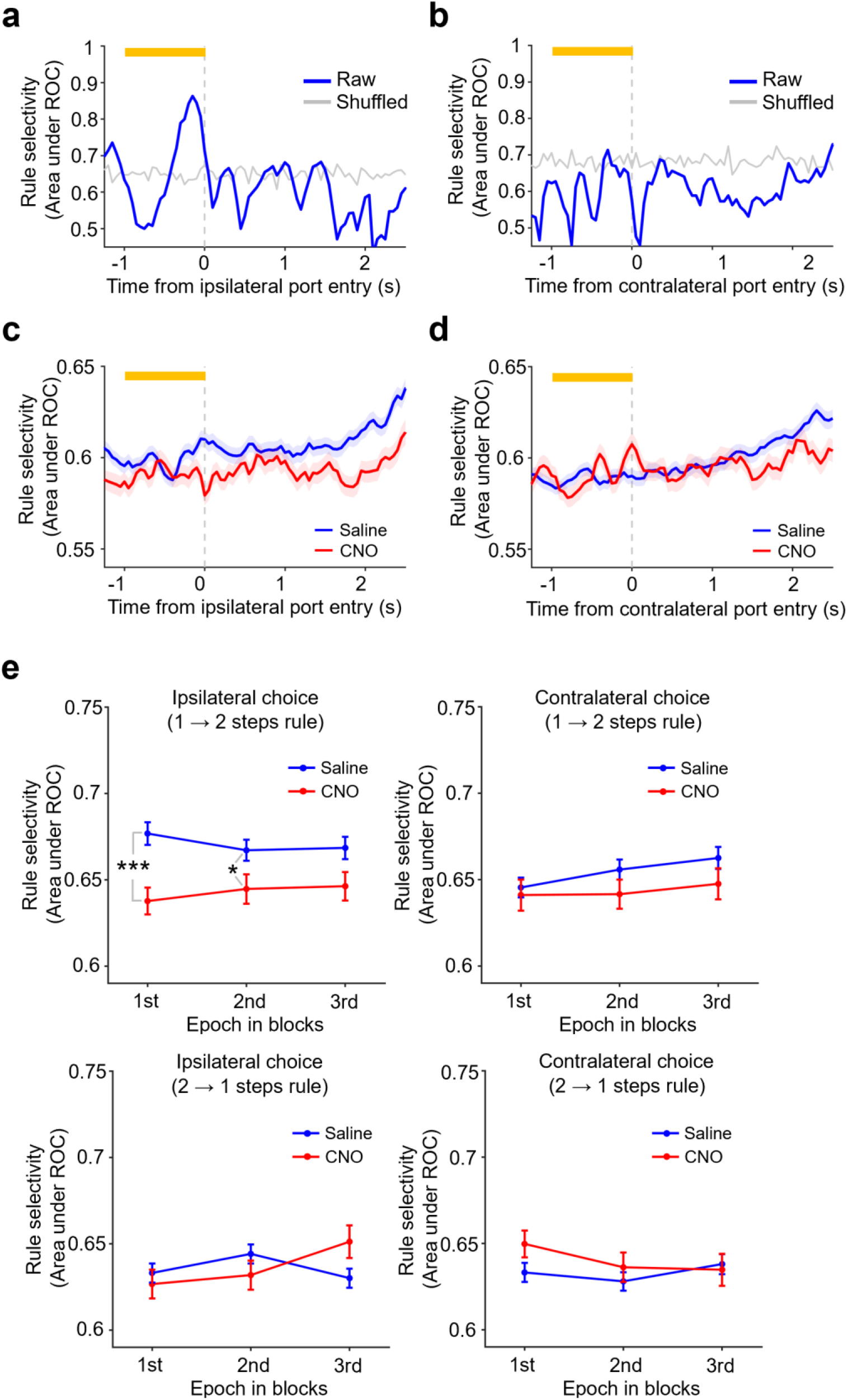
Chemogenetic silencing of ACC decreased rule selectivity in M2 neurons. **a,** Time course of rule selectivity of a representative M2 single-unit (the same unit as presented in Fig. 4b,c) was plotted for trials in ipsilateral condition. Rule selectivity was quantified using an area under ROC curve that was constructed from the distributions of mean firing rates during pre-choice period in trials of Rule Switch blocks of 1 step and 2 steps rule conditions (see main text and Methods for details). Gray line represents a 95% percentile level estimated by the shuffled data in which the area under ROC curve was calculated with rule labels for trials (i.e., 1 step or 2 steps rule conditions) being randomly shuffled. Error bar, s.e.m. **b,** Same as in **a**, but for trials in contralateral condition. **c,** Population-averaged time course of rule selectivity in ipsilateral condition. Blue, saline solution. Red, CNO solution. **d,** Same as in **c**, but for trials in contralateral condition. **e,** Comparison of rule selectivity between CNO and saline conditions and across three epochs in Rule Switch blocks (1st epoch, 1-18th trials; 2nd epoch, 17-36th trials; 3rd epoch, 37-55th trials). See Methods for full details of calculating rule selectivity in each epoch. Briefly, to estimate rule selectivity in each epoch, ROC curves were calculated using distributions of mean firing rates during pre-choice period of trials in the 2nd and 3rd epochs in the preceding block and of mean firing rates during the same period of trials in the 1st, 2nd or 3rd epoch of the subsequent block. A repeated measures two-way ANOVA (with epoch being a within-subject factor) was conducted for ipsilateral condition in blocks following rule switches from 1→2 steps rules. No interaction was found between CNO does and epoch in Ipsilateral choice (1→2 steps rules) condition (*P* > 0.4). A significant main effect of CNO dose was detected (F_1,1833_ = 12.7, *P* = 3.7 x 10^-4^), but not for epoch (*P* > 0.7). Post-hoc comparison using two independent samples *t*-test showed significant differences between saline and CNO solutions in 1st and 2nd epochs. No significant interaction or main effect was detected for other three conditions (i.e., ipsilateral condition in blocks following rule switches from 2→1 steps rule condition, contralateral condition in blocks following rule switches from 1→2 steps rule condition and contralateral condition in blocks following rule switches from 2→1 steps rule condition). ***, *P* = 4.4 x 10^-4^. *, *P* = 0.036. n = 437 and n = 195 single-units for saline and CNO conditions, respectively. Error bar, s.e.m.

### Optogenetic silencing of ACC circuits induced errors in trials immediately following 1 step to 2 steps rule switches

We wanted to determine what aspects of behavioral adjustments upon rule switches require anterior cingulate circuits. To address this question, we optogenetically suppressed ACC excitatory neurons using halorhodopsin (eNpHR3.0) (Fig. 7a). 561 nm laser light delivery decreased spiking activity of ACC neurons (Fig. 7b), which validated *in vivo* neuronal inhibition. We delivered the light for 4 sec immediately following the animals’ incorrect 2nd choices (Fig. 7c) or the animals’ correct 2nd choices (Fig. 7d). In a representative session, light was delivered upon animal’s making incorrect 2nd choices. The animal showed increased error in 2nd choices in trials following light delivery (Fig. 8a). Group results demonstrated that optogenetic suppression after the animal’s erroneous 2nd choices induced such errors in 2nd choices in the trials that followed initial errors (Fig. 8b). This effect was specifically observed in the 1st epoch of Rule Switch blocks but not in other epochs in these blocks or any epoch in the 1st block. Also, the error rate of 1st responses in these animals were unaffected, suggesting that the animals failed to update their sequential choice responses due to ACC inhibition (Fig. 8d). We repeated similar experiments with 4 sec light applied immediately after the animals’ correct 2nd choices (Fig. 7d). Light delivery did not affect 2nd choice performance in any epoch of the 1st or Rule Switch blocks (Fig. 8c). Also, the error rate of 1st responses were unaffected (Fig. 8e). The observed effect of optogenetic silencing after the animals’ incorrect 2nd choice on their task performance (Fig. 8b) could not be explained by heat of light because the same duration of light delivery did not affect the task performance when the light was delivered after the animals’ correct 2nd choices (Fig. 8c). These results indicate that ACC neurons process error feedback information following an erroneous 2nd response and use this information to adjust the animal’s sequential choice responses in subsequent trials.

**Fig. 7.**
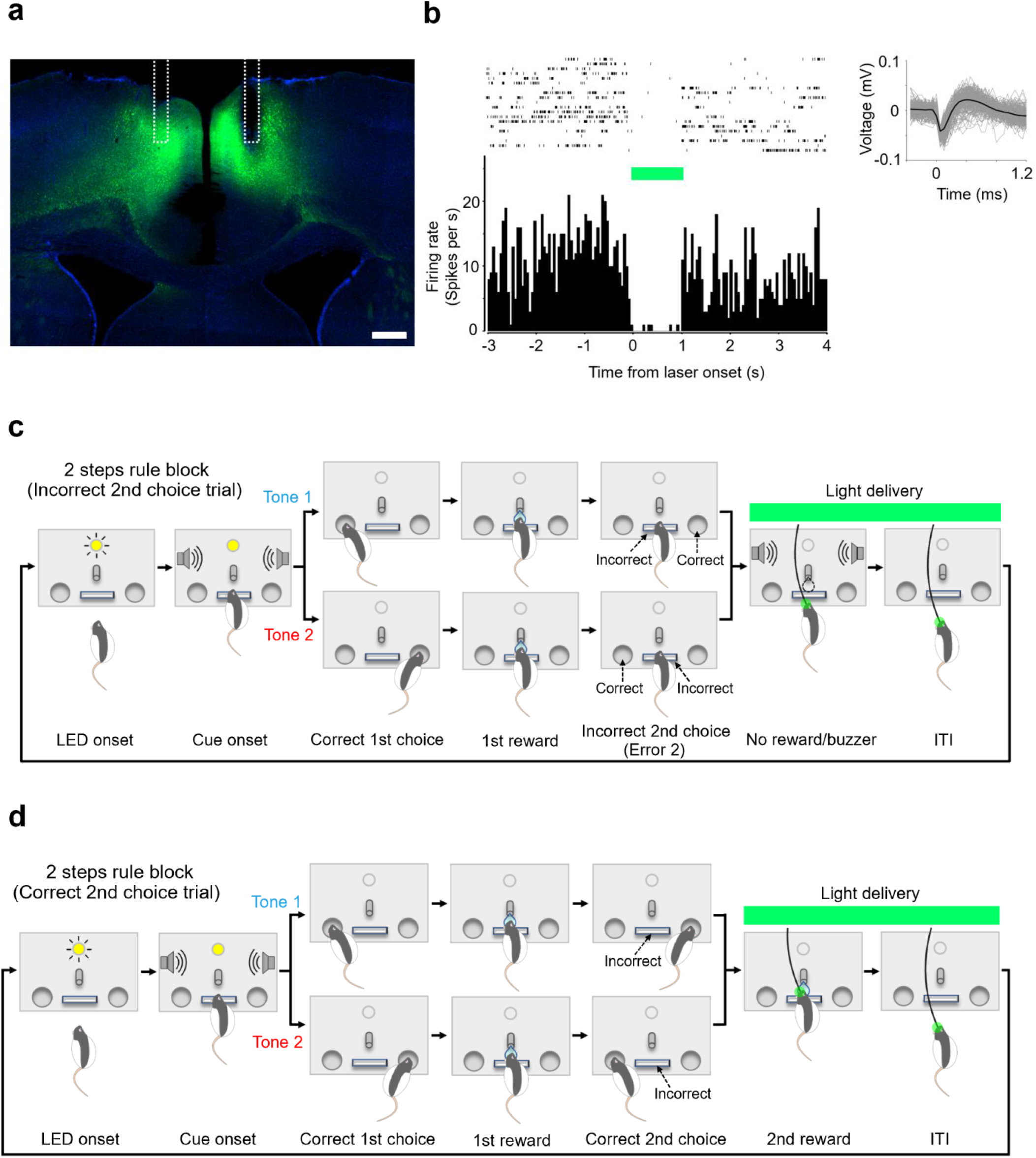
Optogenetic silencing of ACC neurons upon incorrect 2nd choices or correct 2nd choices. **a,** Histological section for halorhodopsin (eNpHR3.0) expression in ACC. White dotted line shows reconstructed positions of fiberoptic implants. Scale bar, 500 μm. **b,** Suppression of spiking activity by 561nm light delivery. Top left, raster plot of a representative single-unit measured in ACC showing spiking activities before, during and after laser light delivery. Top right, example waveforms of the representative single-unit. Bottom left, peri-event time histogram sorted by the timing of light onset. Bin width, 50 ms. **c,** Optogenetic silencing of ACC after animal’s incorrect 2nd choices. Light was delivered for 4sec after an incorrect 2nd choice (i.e., animal’s pushing the center lever before poking the side port opposite to the 1st choice). **d,** Optogenetic silencing of ACC after animal’s correct 2nd choices. Light was delivered for 4sec after a correct 2nd choice (i.e., animal’s poking the side port opposite to the 1st choice).

**Fig. 8.**
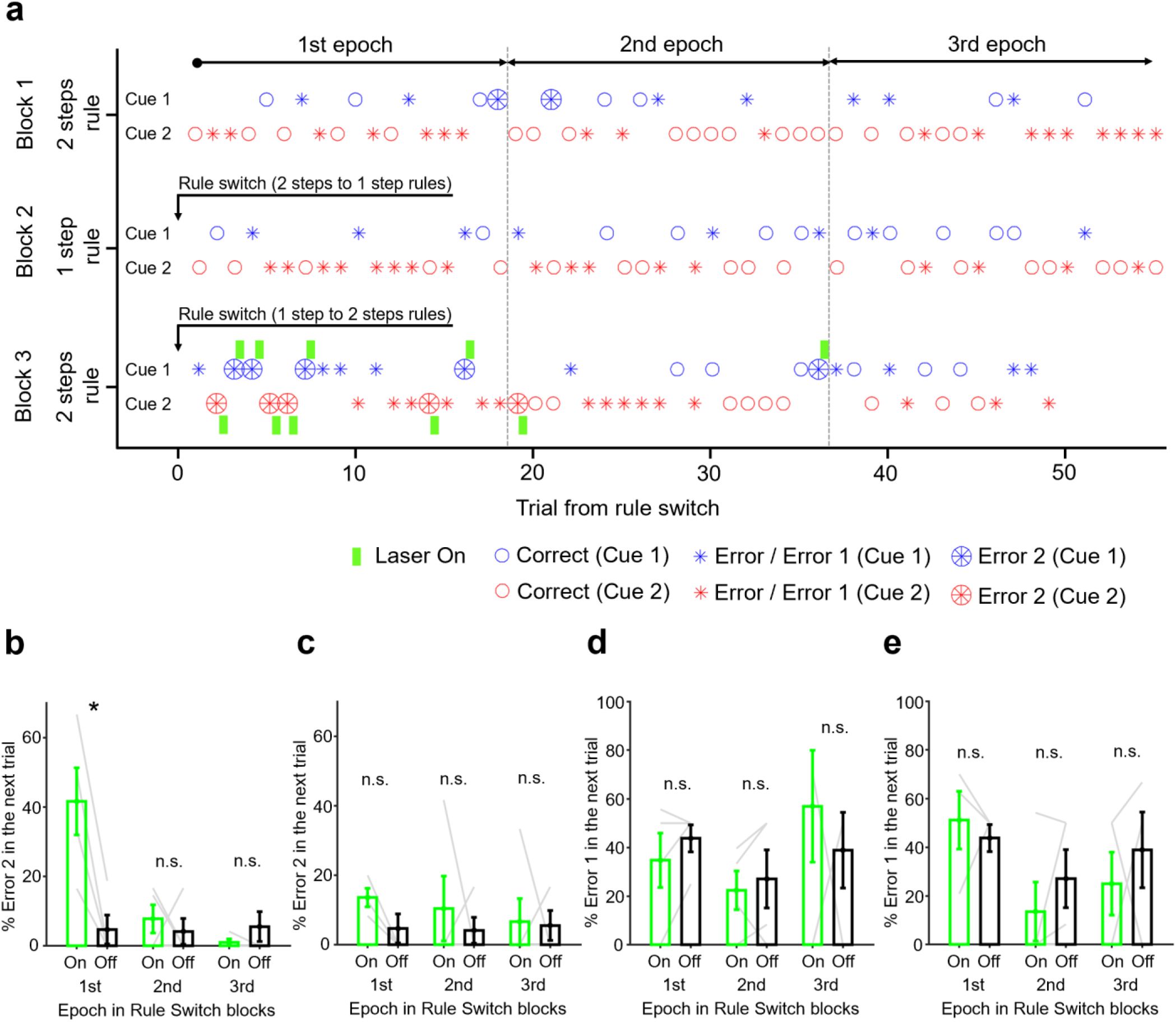
Optogenetic silencing of ACC neurons upon incorrect 2nd choices induced sequential choice errors in the immediately subsequent trials that followed rule switches. **a,** Task performance chart of a representative session in which light was delivered upon animal’s making an incorrect 2nd choice in Rule Switch block (“Block 3”). Note that no light was delivered in the 1st block (“Block 1”) or in any trial of 1 step rule block (“Block 2”). Green bar, light delivery. Small circle, correct trial. Asterisk, incorrect 1st choice (Error 1). Large circle filled with an asterisk, incorrect 2nd choice (Error 2). Blue and red represent trials with two distinct tone cues. Gray dotted lines show borders that separate three epochs in each block (1-18th, 19-36th and 37-55 trials for 1st, 2nd and 3rd epochs, respectively). **b,** 2nd choice performance (%Error2) was plotted for trials that immediately followed an Error 2 trial. “On” and “Off” represent trial conditions in which light was delivered and not delivered. The performance was plotted for each of three epochs separately. *, *P* = 0.0201 (two samples *t*-test, n = 4 rats). Error bar, s.e.m. **c,** Light was delivered for 4 sec after animals’ correct 2nd choices instead of incorrect 2nd choices (see Fig. 7d). % Error 2 was plotted for trials that immediately followed an Error 2 trial. **d**, Same as in **b**, but 1st choice performance (%Error1) was plotted for trials that immediately followed an Error 2 trial in which a light was delivered (“On” condition) upon incorrect 2nd choices or not delivered (“Off” condition). *P* = 0.212, 0.598 and 0.913 for each epoch, respectively. n = 5 rats. **e**, Same as in **c**, but 1st choice performance (% Error1) was plotted for trials that immediately followed an Error 2 trial in which light was delivered (“On” condition) upon correct 2nd choices or not delivered (“Off” condition). *P* = 0.640, 0.503 and 0.587 for each epoch, respectively. n = 5 rats.

## Discussion

Previous studies have suggested that anterior cingulate circuits are recruited when a greater cognitive control is needed for an animal to resolve uncertain situations in which it experiences unexpected errors or conflicts among multiple choice options, needs to use error feedback for future decisions, or obtain new information for updating error likelihood^15–18, 20, 30–32^. Our results suggest that ACC circuits monitor negative outcomes and transmits this information to M2 circuits for reorganizing rule representations for motor outputs. Inhibiting ACC circuits caused M2 neurons to decrease rule selectivity even before animals make first responses. This effect was observed specifically in epochs immediately following rule switches that require animals to discard single step responses and instead to make sequential responses. Interestingly, this effect was observed only in trials during which animals made ipsilateral responses in the first choice followed by contralateral responses. This indicates that ACC circuits reorganize rule representations in M2 neurons that prepare for making second actions even before animals’ make their first choices. This process occurs specifically in trials immediately following rule switches. Given that neurons in premotor and supplemental motor areas can encode planning for several movements ahead^5^, our data suggests that, after rule switches, ACC circuits generate signals that reorganize such neural representations for planning sequential movements in M2 that is crucial in the new rule condition (i.e., representations of second responses).

We observed an effect of chemogenetic silencing of ACC circuits in updating rules from 1 step to 2 steps but not in the opposite direction (i.e., rule switches from 2 steps to 1 step) as measured by non-rewarded second actions. This asymmetry may reflect the asymmetry of our task requirements (i.e., the difference in complexity of choice actions in 1 step and 2 steps -conditions) or may reflect some inherent network characteristics in ACC→M2 circuits such as distinct temporal scales of transitions of population neural activity in M2^33^. A previous study showed that neurons in M2 projecting to dorsolateral striatum regulates the initiation of sequential movements whereas sequence completion depends on activity of striatonigral medium spiny projection neurons in dorsolateral striatum^13^. By analogy, updating rules from 2 steps to 1 step which requires a deletion of the second step response might depend on dorsolateral striatum circuits rather than ACC→M2 circuits. Given that more global brain networks are recruited when an animal makes movements that are not instructed by the behavioral task and are not rewarded^34^, brain networks beyond ACC→M2 circuits (cortical or subcortical) might be necessary to extinguish such non-rewarded movements upon rule switches.

Interestingly, we found no effect of chemogenetic silencing of prelimbic/infralimbic cortex (a phylogenetic homolog of dorsolateral prefrontal cortex or DLPFC in primates) while previous studies have shown that DLPFC also plays a critical role in updating rule representations in the brain^19, 28, 35^. In this study, we minimized the within-trial working memory load while requiring animals to hold rule representations (e.g., “rule memory” or “task set”) across trials in specific rule blocks. The cognitive loads of different time scales (i.e., memory spanning a single trial period is several seconds vs. across rule blocks is ∼10 minutes) might affect how an update of behavioral response would rely on ACC vs. DLPFC circuits^30^. These findings suggest that ACC and DLPFC circuits play distinct roles in behavioral flexibility, which is an exciting direction for future studies.

## Methods

### Rats

All procedures relating to rat care and treatment conformed to the institutional and NIH guidelines. Wild type male Long-Evans rats (> 300 grams) were used (Charles River). Rats were pair-housed during initial behavioral training and then single housed after being injected with viruses or being implanted with electrodes, fiberoptic implants, or cannulae. Rats were kept on a reverse 12 h light/dark cycle, and trained and tested in their dark cycle. Food was available ad libitum, and rats had scheduled access to water for motivating them to work for water reward while monitoring their body weight to ensure they were over 85% of initial weight.

### Behavioral apparatus

Behavior took place in a custom-made chamber (415 mm length, 300 mm width, 500 mm height) inside a sound attenuating cubicle (MED Associates). The cubicle was electromagnetically shielded by copper mesh sheet or nickel/silver fabric in electrophysiology recording experiments. The behavioral setup consisted of a stainless-steel lever at the center (MED Associates) and two ports equipped with infrared photodiodes on the left and right sides of the lever, arranged side-by-side with a center-to-center distance of 65 mm on a stainless-steel wall. An interruption of the infrared beam signaled port entry. A sipper tube was installed on the front wall 25 mm above the center lever and was connected to a water supply that was controlled by a computer-controlled solenoid. In addition, there were two speakers mounted on the side walls (about 150 mm away from the center lever). Timing of presentations of sounds from the speakers (i.e., cue stimulus tones or feedback buzzer sound) and delivery of water rewards were controlled using a multifunction digital input/output board (National Instruments) with custom programs written in C++ and Labview (National Instruments) on a computer running a Windows 10 operating system. Behavioral events were timestamped with a precision of <1 ms.

### Conditional action sequencing (CAS) task

Rats were trained and tested on an auditory cued- two alternative forced choice task as follows. Rats self-initiated each trial with a push on the center lever to receive the tone cue stimulus. After a delay of 10 ms, a tone of either 8 or 12kHz with a sound pressure level of 75 dB was presented in pseudorandom order in each session. The tone was kept on until an animal entered one of the side ports (left or right) as the 1st choice. Choices were rewarded with ∼25 μl water if they poked the correct side port. An error feedback buzzer sound was delivered as a penalty if they poked the incorrect side port followed by an elongated period of inter trial interval (ITI). A feedback buzzer sound accompanied by an elongated ITI was also delivered if animals made a non-rewarded 2nd action (Nr2A), that is, entered the opposite side port after making a correct 1st choice and before initiating the next trial (pushing the center lever after an ITI). In 1 step rule condition, this completed a trial and, after an ITI period (3-4 s following a correct choice trial and 5-6 s following an incorrect choice trial), an LED turned on, signaling the animals to initiate the next trial by pushing the center lever. In the 2 steps rule condition, animals were required to make a 2nd response by entering the side port opposite the one that animals’ chose as the 1st response, instead of pushing the center lever. A correct entry to the side port as a 2nd response was rewarded with 25 μl water. An error feedback buzzer sound was delivered if they poked the incorrect side port.

1 step and 2 steps rule conditions were switched in every 55 trials (with exceptions of 2 sessions in which rules were switched in every 40 trials) in a block-wise manner. In some sessions, behavior experiments started from the 2 steps rule condition (1st block) and then proceeded to the 1 step rule condition (2nd block) and so forth (see Extended Data Fig. 1a). In other sessions, experiments started from the 1 step rule condition in the 1st block so that animals’ task performance in 1st block and Rule Switch blocks in 1 step rule condition could be compared. There was no explicit cue signaling the animals that there was a rule change between blocks. Therefore, animals needed to change their responses in the 2nd choice (either to poke the opposite side port or pushing the center lever to initiate the next trial without poking the side port) solely based on the omission of reward after the 2nd choice and the delivery of a feedback buzzer sound.

### Training

Rats were trained over the course of 8-12 weeks, with progressive introduction of each aspect of the task as follows. After handling and habituation sessions in the behavior box (2-3 days), rats were first trained to either push the lever or enter a side port (left or right) to receive a water reward. Next, rats were trained to perform a contingency task in which a tone sound (4kHz) was delivered when the animal pushed the center lever and then the animal was required to poke either left or right side port to receive a reward (typically required several sessions). Then rats were trained on auditory discrimination in which they were required to choose left or right side ports depending on the tone cue stimulus (either 8 or 12kHz). Once they reached over 70% correct performance for two consecutive days, they were trained with the conditional action sequencing (CAS) task without a rule change between the 1 step and 2 steps rule conditions, requiring animals to make 2 step responses throughout the session. Once they reached over 70% correct performance for two consecutive days, they were trained on the conditional action sequencing (CAS) task with rule changes between the 1 step and 2 steps rule conditions once every 70- 100 trials per block. The length of a block was gradually shortened to 55 trials to complete the task training phase. In many training sessions and in some testing sessions, we adjusted the proportion of two trial types (i.e., tone cues) to abate animals’ choice bias to specific side ports. Such adjustments were typically limited to a ratio of less than 2.

### Surgery

All surgeries were performed under isoflurane anesthesia (1.0-2.0%) using standard stereotactic technique. Following an intraperitoneal administration of a cocktail solution of ketamine (80 mg kg^-1^) and xylazine (8 mg kg^-1^), rats were placed in an isoflurane induction chamber for 5-10 minutes. Then rats were moved to a stereotactic frame and their nose was placed in a cone, which provided 1.5-2% continuous isoflurane flow. After verifying surgical levels of anesthesia with pinch tests and eye blink tests, rats were secured in non-rupture ear bars (Kopf Instruments). The concentration of isoflurane was maintained at 0.75-1.5% throughout the surgery. Slow release buprenorphine (1mg kg^-1^) and a Ringer’s solution were administered in the middle and at the end of the surgery, respectively. Rats remained on a heating pad until they made a full recovery from anesthesia after surgery. Supplemental nutrient gels were supplied to rats after they were returned to their home cages. Rats were monitored during their recovery from surgery for at least 4 days before restarting experiments.

### Viral injections

For chemogenetics experiments, we used AAV2/5-CaMKIIa-hM4Di-mCherry and AAV2/5-CamKIIa- mCherry viruses. The plasmids were obtained from Addgene (Plasmid#50477 and Plasmid#114469) and packaged by Vigene after in house plasmid preparation. For optogenetics experiments, we used AAV2/9- CamKIIa-eNpHR3.0-eYFP virus. The plasmid was obtained from Addgene (Plasmid#26971) and packaged by Vigene after in house plasmid preparation. The viral titers were 3.3 x 10^14^ genomic copy (GC) ml^-1^ for AAV2/5-CamKIIa-hM4Di-mCherry, 1.9 x 10^13^ GC ml^-1^ for AAV2/5-CamKIIa-mCherry and 1.3 x 10^13^ GC ml^-1^ for AAV2/9-CamKIIa-eNpHR3.0-eYFP virus. For viruses used in viral tracing experiments, see the following section for viral tracing experiments.

Each animal underwent bilateral craniotomies using a 1/4 size drill bit. The virus solutions with a volume of 1 μl were injected using a mineral oil-filled glass micropipette joined by a microelectrode holder to a 10 µl Hamilton micro syringe. A micro syringe pump was used to control the speed of virus injections (2 nl min^-1^). The micropipette was slowly lowered to the target site and remained for 5 minutes before starting injections (ML ±1.3 mm, AP +2.0 mm, DV -0.9 mm for M2; ML ± 0.5 mm, AP -1.0 mm, DV - 1.4 mm for anterior cingulate cortex; ML ± 0.6 mm, AP +3.0 mm, DV -3.0 mm for prelimbic/infralimbic cortex; ML ± 2.0 mm, AP -2.2 mm, DV -6.0 mm for ventral thalamic nuclei). After injections, the micropipette stayed for 10 min before it was withdrawn.

### Viral tracing experiments

For exploring brain regions projecting to M2 in the rat, we used a genetically engineered rabies virus^25^. We first unilaterally injected a cocktail solution of pENN.AAV.CaMKII.0.4.Cre.SV40 (Addgene#105558-AAV9) and AAV2/rh8-synP-DIO-sTpEpB-WPRE-bGH with a mixed ration of 1:1 at M2 (1 μl, ML ±1.3 mm, AP +2.0 mm, DV -0.9 mm)^36^. 1-2 weeks later, RVΔG-4mCherry (EnvA) (1 μl) was injected at the same coordinates^37^. Viral titers for the injected solution were 1.05 x 10^13^ GC ml^-1^ and 1.15 x 10^12^ GC ml^-1^ for pENN.AAV.CaMKII.0.4.Cre.SV40 and AAV2/rh8-synP-DIO-sTpEpB-WPRE-bGH, respectively, and 1.7 x 10^10^ GC ml^-1^ for RVΔG-4mCherry (EnvA). One to two weeks (typically ∼7 days) after the injection of rabies virus, the animal was perfused, bran extracted, sectioned, immuno- stained and imaged.

To validate projections from ACC to M2, we injected AAVretro-pmSyn1-EBFP-Cre (Addgene#51507-AAVrg) at M2 (1 μl, ML -1.3 mm, AP +2.0 mm, DV -0.9 mm) and AAV5-hSyn-DIO- hM4Di-mCherry (Addgene#44362) at anterior cingulate cortex (ML -0.5 mm, AP -1.0 mm, DV -1.4 mm). The viral titers were 1.1 x 10^13^ GC ml^-1^ for AAVretro-pmSyn1-EBFP-Cre and 4.7x10^12^ GC ml^-1^for AAV5- hSyn-DIO-hM4Di-mCherry.

### Immunohistochemistry

Rats were deeply anesthetized using sodium pentobarbital and then transcardially perfused with saline and 4% paraformaldehyde (PFA). Brains were extracted and incubated in 4% PFA at room temperature overnight. Brains were transferred to PBS, and 50 μm coronal slices were prepared using a vibratome. For immunostaining, each slice was placed in PBS + 0.1% Triton X-100 (PBS-T), with 10% normal goat serum for 1 hr and then incubated with primary antibody at 4℃ for 12 hr. Slices then underwent three wash steps for 10 min each in PBS-T, followed by 2 hr incubation with secondary antibody. After three more wash steps of 10 min each in PBS-T, slices were transferred to DAPI solution (5 μg ml^-1^ in PBS), incubated for 30 min in room temperature and then mounted on microscope slides. Antibodies used for staining were as follows: to stain for hM4Di-mCherry or mCherry alone, slices were incubated with primary rabbit anti-RFP (1:1000, Rockland) and visualized using anti-rabbit Alexa 555 or Alexa 568 (1:200). To stain for eNpHR3.0-eYFP, slices were incubated with primary chicken anti-GFP (1:1000, Life Technologies) and visualized using anti-rabbit Alexa 488 (1:200). Immuno-stained slices were imaged using an epifluorescence (Zeiss Imager.Z2) or a confocal microscope (Zeiss LSM700) with 5X or 10X objective lenses. Intensity of each fluorescence channel in imaging data were adjusted using ZEN blue software.

### Chemogenetic experiments

After the injection of inhibitory DREADD virus (AAV5-CamKIIa-hM4Di-mCherry), we tested rats’ behavioral performance in the CAS task. 10 or 20 mg kg^-1^ solution of clozapine-N-oxide (CNO; Sigma, C0832-5MG) was prepared by first dissolving CNO in dimethyl sulfoxide (DMSO; Sigma, 34869) followed by adding saline solution (final concentration of DMSO were 5% and 1% for intraperitoneal injection and for cannula infusion experiments, respectively). CNO solutions were intraperitoneally (*i.p.*) injected 35-40 min before starting behavioral testing. In some sessions, we started behavior experiments 60 min after the *i.p.* injection of CNO solution to examine whether the duration from the CNO administration to the start time of behavioral testing could affect the animal’s performance. In saline control conditions, a vehicle solution (5% DMSO in 0.9% saline) was intraperitoneally injected 35-40 minutes before behavioral sessions.

For locally infusing CNO solution or its control solution in M2 in rats infected with inhibitory DREADD virus in ACC, we implanted 26 gauge dual guide cannula (Plastics One) targeting bilateral M2 (ML ±1.3 mm, AP +2.0 mm). Rats were placed under a light non-surgical 1-1.5 % isoflurane anesthesia and the CNO solution (0.5μl, 1 μg μl^-1^) was bilaterally infused by through 33 gauge dual internal cannula (Plastics One) at a speed of 0.2 μl min^-1^. After waiting for 4 min after infusions, the infusion cannula were removed. Rats were kept in their home cage for 30 minutes until we started behavioral experiments.

For testing if an intraperitoneal administration of CNO could suppress neural activity in ACC, we measured multiunit spiking activity in ACC in two rats infected with the inhibitory DREADD virus and implanted with silicon probes (Neuronexus). We first intraperitoneally injected a saline control solution (5% DMSO in 0.9% saline) immediately before starting neural activity measurements and continued the electrophysiological recording for 60 min. Then animals were transferred to the isoflurane chamber. After intraperitoneal injection of the CNO solution (10 or 20 mg kg^-1^ in 5% DMSO saline solution) under a light anesthesia, we resumed neural activity measurements. Continuous voltage signals were recorded in hard disc for offline data analysis. To obtain multiunit spike timestamps in offline analysis, the continuous voltage signals were high-pass filtered (400kHz) digitally, and multiunit spikes were detected by thresholding the continuous signals at 4 SDs above the baseline level^38^.

### Electrophysiology in task behaving rats

The neural activity data was obtained from six rats in which Utah array electrodes with 4 x 8 matrix shanks (platinum or iridium-oxide tips; electrode length of 0.5 mm or 1 mm; electrode spacing of 0.4 mm) (Blackrock Microsystems) were implanted in M2 targeting an area covering the Bregma coordinate of ML 1.3 mm, AP +2.0 mm in either left or right hemispheres (left hemisphere in two animals and right hemisphere in three animals). This location was chosen because it was the center of the distribution of stimulation sites that resulted in contralateral orienting movement and neurons related to orienting responses were recorded in previous studies^8,39^. We also confirmed that delivering electrical stimulations (20 s^-1^ bipolar injections of 30-60 μA current) at around this coordinate elicited contractions of shoulders or limbs of the rats.

For implant surgery, animals were anesthetized in the isoflurane chamber and placed in the stereotaxic frame. After applying eye ointment and washing the incision sites with betadine and ethanol, an incision was created over the scalp and connective tissue was removed. The skull was drilled for attaching bone screws and for implanting the array electrodes in M2. After installing bone screws, we performed a durotomy and slowly inserted the array electrodes at M2 using a manipulator and a mounting probe by applying a vacuum to steady the array. Dental cement was applied to secure the cable to the screws and the craniotomy was filled with a surgical silicone adhesive (Kwik-sil). The vacuum was turned off and the array was further secured using dental cement.

All recordings were conducted after the rats were fully recovered (at least seven days after surgery). The ground was taken from one of the skull screws typically above the cerebellum. The reference channel was chosen from one of the skull screws or one of the recording channels in which no clear spiking activity was observed. Data was recorded using a unity gain amplifier forwarded, filtered (600-8000Hz, FIR filter) and stored in the Digital Lynx SX System (Neuralynx) for offline data analysis.

### Optogenetic manipulation in behavior experiments

Dual fiberoptic cannulae (dual 200 µm core, NA = 0.22, Doric Lenses) was implanted in anterior cingulate cortex in task trained rats under isoflurane anesthesia (see surgery section for general procedures). A yellow-green laser (Opto Engine LLC, 561 nm) with a fiber optic patch cable (dual 200 µm core, NA = 0.22, Doric Lenses) was installed on the behavioral chamber. A TTL pulse was delivered from the behavior system through an interface board (National Instrument) that determined the laser timing for optogenetic intervention. The output power of the laser to the bilateral fiberoptic cannula was calibrated to 15mW per channel for the yellow-green laser with the implanted optical fiber attached. This power was determined by acute optogenetic experiments (see below).

An optrode consisting of a tungsten electrode (0.5 MΩ or 1MΩ; FHC Inc.) attached to an optical fiber (200 μm core diameter; Doric Lenses) with the tip of the fiber extending beyond the tip of the electrode by 200-300 μm was used for simultaneous optical stimulation and extracellular recordings. The optrode was slowly lowered to cingulate cortex where the viruses were injected. The optical fiber was connected to a green laser (561 nm, 200 mW; MBL F561, OptoEngine) and controlled by a beam shutter and a shutter controller. The power intensity of light emitted from the optrode was calibrated to 13-16 mW as measured with an optical power/energy meter, which is consistent with the power intensity used in behavior experiments. At each depth of the optrode, 1sec light pulses were delivered at 0.2 s^-1^ and neuronal activity was collected for 10-20 sweeps for each recording session. Continuous voltage data were monitored online using an oscilloscope and a sound speaker, fed into a preamplifier, transferred to an interface board and saved in the hard disc for offline analysis. Rats were sacrificed, and brains were collected and sectioned for histological confirmation of recorded sites.

### Data analysis

Matlab and R were used for data analysis (see the following section for details). No statistical analysis was conducted to pre-determine sample sizes of each experiment. Statistics were run two-sided except mentioned otherwise.

### Statistics

In testing statistical significance of the effect of CNO dose or epoch in Rule Switch blocks or their interaction in task performance (Figs 2g-k and 4d-g), we conducted repeated measures two-way ANOVA with both CNO dose and epoch as within-subject factors by using “aov” function provided in R. In testing statistical significance of the effect of CNO dose or epoch in Rule Switch blocks or their interaction in rule selectivity (Fig. 6e), we conducted a repeated measures two-way ANOVA with CNO dose and epoch as between-subject and within-subject factors, respectively, by using “Anova” function in “car” library in R. All other statistical tests were conducted using Matlab.

### Chemegenetics behavior data analysis

We used a total of 29 rats for chemogenetics experiments; 15 rats were injected with the inhibitory DREADD virus (AAV2/5-CaMKIIa-hM4Di-mCherry) in ACC, 5 rats were injected with the mCherry control virus (AAV2/5-CaMKIIa-mCherry) in ACC, 4 and 5 rats were injected with the inhibitory DREADD virus in prelimbic/infralimbic cortex and ventral thalamic nuclei, respectively. Among the 15 rats that were injected the inhibitory DREADD virus in ACC, 11 rats were used for experiments intraperitoneally administering CNO solutions (n = 3 rats for only 20 mg kg^-1^ dosage, n = 3 rats for only 10 mg kg^-1^ dosage and n = 5 rats for both 20 mg kg^-1^ and 10 mg kg^-1^ dosage) while 5 rats were used for experiments locally infusing CNO solutions with a dose of 1 μg μl^-1^ (one rat was used for both *i.p.* injection experiment and cannula infusion experiment). Two animals (three sessions) were also tested with an *i.p.* injection of CNO with a dose of 40mg kg^-1^ but the data were not included in the group analysis. Also, one rat was pilot-tested with local infusion of CNO solutions with a dose of 0.4 μg μl^-1^ and 4 μg μl^-1^ but these data were not included in group analysis.

To compare animals’ task performance between the 1st block in the session and the following Rule Switch blocks (either 1 step or 2 steps rule block), sessions were not included in the group analysis if they don’t have Rule Switch blocks (i.e., if animals stopped task behaviors before the task entered in Rule Switch blocks). Also, to compare animals’ task performance across three epochs within a block (either 1 step or 2 steps rule -block), sessions were not included in the group analysis if they have less than 10 trials in the 3rd epoch of at least one Rule Switch block. To compare the task performance in the first, middle and third sections in a block, a block was subdivided in three epochs: trials no. 1-18 (1st), 19-36 (2nd) and 37- 55 (3rd) for sessions with rule switches in every 55 trials. Similarly, in two sessions in which the rule was switched in every 40 trials, a block was subdivided in three epochs: trials no. 1-13 (1st), 14-26 (2nd) and 27- 40 (3rd).

Animal’s task performance was measured by percent error rate of the choice in 1 step rule block (“%Error”), percent error rate of the 1st choice (“%Error 1”) and that of the 2nd choice (“%Error 2”) in 2 steps rule block. %Error and %Error 1 were used to probe animal’s capacity to make auditory discrimination as well as its capacity to make choice response actions. %Error 2 was used to probe animal’s capacity to make correct sequential choice responses. Note that animal can choose to push the center lever for initiating the next trial instead of poking a side port opposite to the one that the animal chose in its 1st response. Such response (i.e., pushing the center lever) was counted as an error response (i.e., %Error 2) in 2 steps rule condition, but was not counted as an error in 1 step block. We used %Error 2 to probe animal’s capacity to adjust its choice responses following rule changes from 1 step to 2 steps conditions (see Extended Data Fig. 1d, right panel). We also measured the frequency of animal’s making a 2nd choice response in 1 step rule block. Such response (referred to as “non-rewarded 2nd action” or Nr2A) was not rewarded and was penalized by a presentation of feedback buzzer sound and by the prolonged ITI duration (see Extended Data Fig. 1d, left panel). Frequency of Nr2A is expected to decrease in 1 step rule block as rats adjusted its responses after the rule switches from 2 steps to 1 step conditions. We used Nr2A to probe animal’s capacity to adjust its choice responses following rule changes from 2 steps to 1 step conditions.

In group analysis, percent error rate for two trial types (i.e., trials with tone cue 1 and those with tone cue 2) were separately calculated and then averaged. Response time for the 1st choice response (RT1) as presented in Extended Data Fig. 4e,f was calculated as the difference in the timing of center lever entry and first choice port entry (Choice1). Similarly, response time for the 2nd choice response (RT2) was calculated as the difference in the timing of second choice port entry (Choice2) and first choice port entry (Choice1).

### Electrophysiology data analysis (chronic recordings from task behaving rats)

In group analysis, as in chemogenetics behavior data analysis, sessions with a total trial number of less than 10 trials in the 3rd epoch of Rule Switch block were not included in the analysis. Trials with outlier RT1 (cut off = 3 sec) were removed from electrophysiology data analysis (also see the following section. In all analyzed sessions, the proportion of trials with outlier RT1 was less than 2% of the total trials of the session).

Single-units were isolated by spike sorting based on peak or valley and/or principal components of the voltage-thresholded waveforms using the Offline Sorter software (Plexon). Only a unit with a refractory period (> 2 ms) in the auto-correlogram was accepted as single-units^38, 40^. We analyzed neural data collected in 43 sessions from 5 rats: 29 sessions with saline control (n = 5 rats, 2-9 sessions for each), 14 sessions with an *i.p.* injection of CNO solution with a dose of 20 mg kg^-1^ (n = 5 rats, 1-4 sessions for each). From these session data, a total of 900 single-units were isolated: 594 units and 306 units for saline and CNO conditions, respectively. All these units were included in the group analysis of mean firing rate (Fig. 5). Spike timestamps of a single-unit in each task trial were smoothed using a gaussian kernel (σ = 60 ms) and peri-event time histograms (PETHs) were constructed with a bin width of 50 ms.

A single-unit was included in the group analysis of rule selectivity (Fig. 6) if it showed mean firing rate of at least 3 spikes s^-1^ in the 1sec window immediately before animal’s making the choice in either cue stimulus condition in 1 step rule condition or before animal’s making the 1st choice in either cue stimulus condition in 2 steps rule condition (437 units and 195 units for saline and CNO conditions, respectively).

To quantify rule selectivity of neural responses, we used receiver operating characteristic (ROC) analysis^27–29^. An ROC analysis measures the degree of overlap between two response distributions. For each M2 single-unit the preferred and non-preferred rule conditions were compared, given two distributions, P and N respectively, of neuronal activity. For example, for some neurons (e.g., a representative neuron presented in Fig. 5b,c), these distributions were the neurons’ firing rates during the 2 steps rule was in effect in comparison to the 1 step rule. An ROC curve was then generated by taking each observed firing rate of the neuron and plotting the proportion of P that exceeded the value of that observation against the proportion of N that exceeded the value of that observation. The area under the ROC curve was then calculated. A value of 0.5 would indicate that the two distributions were completely overlapped, and thus the neuron is not selective to the rules. A value of 1.0, on the other hand, would indicate that the two distributions were completely separated (i.e., every value drawn from N is exceeded by the entirety of P, whereas none of the values of P is exceeded by any of the values in N) and so the neuron is very selective. This method of analysis has the advantage that it is independent of the firing rate, and so can be used to compare neurons with different baseline firing rates and dynamic ranges^29^. It is also nonparametric and so does not require the distributions to be Gaussian.

We calculated the mean firing rate of each single-unit during a 1 sec period immediately before animals made their 1st choices in each trial of either 1 step or 2 steps rule conditions (pre-choice period). We then calculated the area under ROC curve (auROC) using these firing rates. To examine how rule selectivity changes throughout the Rule Switch blocks, a block was subdivided in three epochs: trials no. 1-18 (1st epoch), 19-36 (2nd epoch) and 37- 55 (3rd epoch) for sessions with rule switches in every 55 trials. Similarly, in two sessions in which the rule was switched in every 40 trials, a block was subdivided in three epochs: trials no. 1-13 (1st epoch), 14-26 (2nd epoch) and 27- 40 (3rd epoch).

Then ROC analysis was conducted for each epoch separately. For example, to calculate rule selectivity in the 1st epoch of a block following rule switches from 1→2 steps rule conditions, ROC curves were calculated using distributions of mean firing rates during pre-choice period of trials in the 2nd and 3rd epochs in the preceding 1 step rule block and of mean firing rates during pre-choice period of trials in the 1st epoch of subsequent 2 steps rule block. Similarly, rule selectivity in the 2nd (or 3rd) epoch of a block following rule switches from 1→2 steps rule conditions were calculated using distributions of mean firing rates during pre-choice period of trials in the 2nd and 3rd epochs in the preceding 1 step rule block and of mean firing rates during pre-choice period of trials in the 2nd (or 3rd) epoch of subsequent 2 steps rule block. Similarly, rule selectivity in each of three epochs of a block following rule switches from 2→1 steps rule conditions were calculated using distributions of mean firing rates during pre-choice period of trials in the 2nd and 3rd epochs in the preceding 2 steps rule block and of mean firing rates during pre-choice period of trials in each of three epochs of subsequent 1 step rule block.

To obtain time courses of rule selectivity, we calculated mean firing rates of each neuron using a sliding window with 250 ms width and a step size of 50 ms throughout a trial and auROC curve was calculated for each time point^28^. To test statistical significance of rule selectivity, we used a bootstrap analysis and repeated the ROC analysis 200 times to obtain 95% percentile threshold in which we assigned the rule condition (i.e., 1 step or 2 steps) at random to each trial and calculated the auROC. In group analysis, trial conditions were grouped in “ipsilateral” and “contralateral” -conditions depending on the hemisphere implanted with the electrode array in each animal: in “ipsilateral” trial conditions, animals were required to first choose the ipsilateral side port and then to the contralateral side port as its 2nd choice (i.e., trials requiring the animal to select the right port as its 1st choice if the animal was implanted with the array electrode in the right hemisphere, and vice versa) while, in “contralateral” trial conditions, they were required to select the contralateral side port as its 1st choice and then to the ipsilateral side port as its 2nd choice (i.e., trials requiring the animal to go to the left port if the animal was implanted with the array electrode in the right hemisphere, and vice versa) (see Fig. 5a).

### Optogenetics data analysis

561 nm laser light was delivered upon animal’s making an incorrect 2nd choice (Fig. 7c) or upon a correct 2nd choice (Fig. 7d). We quantified the percent error rate of 1st choice (% Error 1) or 2nd choice (% Error 2) in trials immediately following trials in which a laser light was delivered upon animal’s committing an incorrect 2nd choice (Fig. 8b,d) or making a correct 2nd choice (Fig. 8c,e). As in the data analysis in chemogenetics data and electrophysiology data, for comparing the task performance in trials immediately following rule switches and trial in later part of the block, trials were categorized in three groups according to their positions in the block: trial no. 1-18 (1st epoch), 19-36 (2nd epoch) and 37- 55 (3rd epoch).

To examine if 561 nm laser light delivery silences neural activities in ACC in rats infected with AAV2/9-CaMKIIa-eNpHR3.0-eYFP virus, we offline analyzed the firing rate of spiking activities of isolated single-units in anterior cingulate cortex that were recorded during optogenetic experiments in anesthetized condition. Continuous voltage traces were high pass filtered with a Butterworth filter and then thresholded typically at around -50 to -80 μV using Offline Sorter software (Plexon). Spike rasters and PETHs were plotted using Neuro Explorer software (Plexon) (see Fig. 7a,b).

## Acknowledgements

We thank J. Derwin, J. Martin, S.Y. Huang, M. Ragion for help with experiments; K. Rockland, J. Yamamoto, A. Bari for experimental advices; M. Pignatelli, Q. Ferry, A. Aqrabawi, K. Flick for comments on the manuscript and all the members of the Tonegawa lab for their support. This work was supported by the RIKEN Center for Brain Science, the Howard Hughes Medical Institute, the JPB Foundation (to S.T.), and the Human Frontier Science Program fellowship (to D.T.).

## Author contributions

D.T., D.R., S.M., T.K. and S.T. designed the experiments. D.T. and C.L. conducted immunohistochemistry and image data acquisition. H.A.S. and I.R.W. constructed rabies virus. D.T. collected and analyzed behavior, electrophysiology and circuit tracing data. D.T. and S.T.: wrote the paper with input from all authors.

## Corresponding authors

Correspondence to Daigo Takeuchi or Susumu Tonegawa.

## Data availability

The data that support the findings of this study are available from corresponding authors upon reasonable request.

## Code availability

The program codes that support the findings of this study are available from corresponding authors upon reasonable request.

---

**Extended Data Fig. 1.**
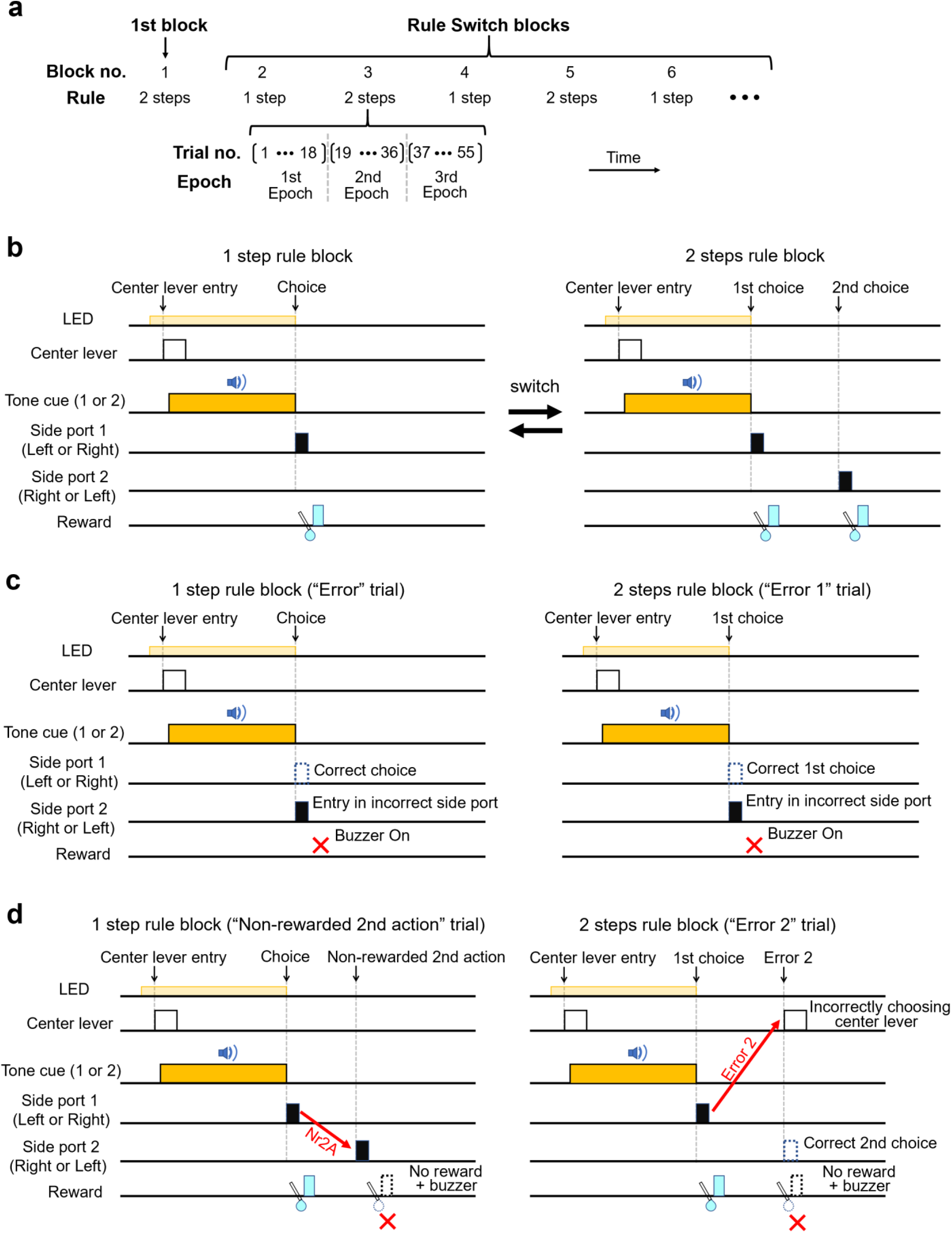
Behavioral task structure (see Fig. 1 main text). **a,** Block structure of the conditional action sequencing task (CAS task). In every 55 trials, task rules were switched between 1 step and 2 steps rule conditions. In some sessions, experiments started with 2 steps rule block (as depicted in the Figure) while, in other sessions, they started with 1 step rule block. All the blocks excluding the first block (“1st block”) were grouped and referred to as “Rule Switch blocks” because there were rule switches preceding these blocks. To compare animals’ task performance in each block, a block was divided in three epochs consisting of 18 or 19 trials (1-18th trial, 19-36th trial and 37-55th trial for 1st, 2nd and 3rd epochs, respectively). **b,** Trial structure of the CAS task in 1 step rule block (left) and in 2 steps rule block (right). LED onset signals the end of inter trial interval (ITI) and rats can start a new trial by pushing the center lever. A tone cue stimulus (either 8kHz or 12kHz frequency tone) was presented when rats triggered the center lever. In 1 step rule condition, rats were required to choose left port or right port depending on the tone cue. If they poked a correct side port (“correct choice”), a water reward was delivered and the next trial could be initiated after an ITI. In 2 steps rule condition, similar to 1 step rule, rats were required to first choose left or right side port depending on the tone cue and, if they poked a correct side port (“correct 1st choice”), a water reward is delivered. Rats received a 2nd reward if they poked the opposite side port as their 2nd choices. Note that the mapping between two cue tones and 1st choice responses was kept the same between 1 step and 2 steps rule conditions while number of response steps were switched between these two rule conditions. **c, “**Error” trial in 1 step rule condition and “Error 1” trial in 2 steps rule condition. In 1 step rule condition, if rats poked an incorrect side port, they received no water reward and instead a buzzer sound was presented with an elongated intertrial interval being imposed before the next trial. Similarly, in 2 steps rule condition, if rats poked an incorrect side port as their 1st choice, they received no water reward and instead a buzzer sound was presented with an elongated intertrial interval being imposed before the next trial. Once rats made an incorrect 1st choice, no reward was delivered even if they entered the opposite side port. **d,** Left, “Non-rewarded 2nd action (Nr2A)” in 1 step rule condition. In 1 step rule condition, no 2nd water reward was delivered when rats poked the opposite side port after making a correct choice and receiving the 1st reward, thus we referred to such response action as “non-rewarded 2nd action (Nr2A).” We used the number of occurrences of Nr2A per trial as an operational measure to quantify animal’s ability to adapt to rule switches from 2 steps to 1 step rule conditions. Right, “Error 2” trial in 2 steps rule condition. In 2 steps rule condition, no reward was delivered and instead a buzzer sound was delivered when rats pushed the center lever after making correct 1st choices and before poking the opposite side port. We referred to this response as “incorrect 2nd choice” or “Error 2”. The percentage of occurrence of this error was referred to as “%Error 2” and was used as an operational measure to quantify animal’s ability to adapt to rule switches from 1 step to 2 rule conditions.

**Extended Data Fig. 2.**
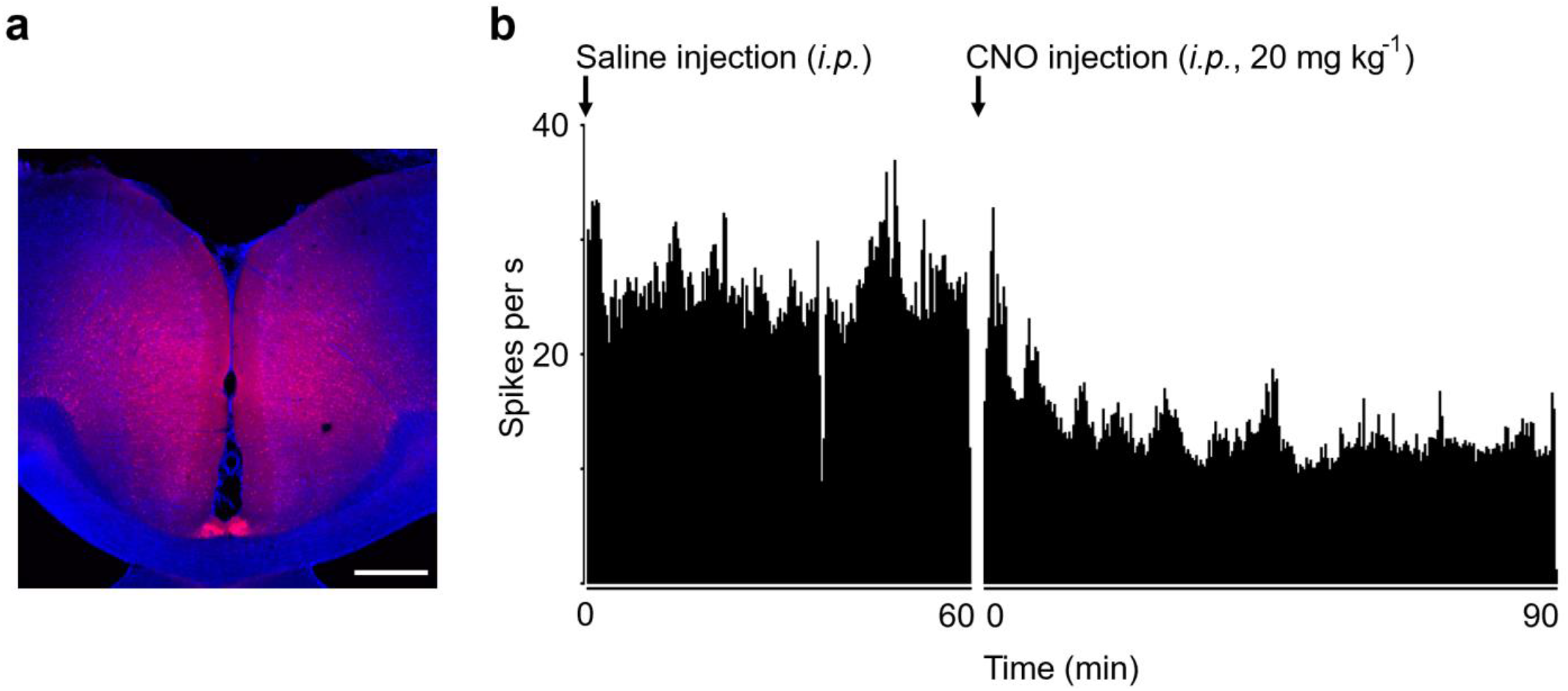
Intraperitoneal injection of clozapine-N-oxide solution suppressed spiking activities in anterior cingulate cortex expressing inhibitory DREADD virus (see Fig. 2 main text). **a,** Inhibitory DREADD virus expression in anterior cingulate cortex (ACC). AAV5-CaMKIIa-hM4Di- mCherry virus was bilaterally injected in cingulate cortex (1000 nl for each hemisphere). Scale bar, 500 μm. **b,** Time histogram of multiunit firing rate measured in ACC expressing inhibitory DREADD virus. First, multiunit spiking activities were measured for 60 minutes after an *i.p.* injection of saline solution. Then, immediately after an *i.p.* injection of clozapine-N-oxide (CNO) solution (20 mg/kg), multiunit activity measurement was resumed that lasted for another 90 minutes. Spiking activity decreased after CNO injection and this effect lasted for at least 60 minutes after having reached a plateau level (∼30 minutes after an administration of CNO).

**Extended Data Fig. 3.**
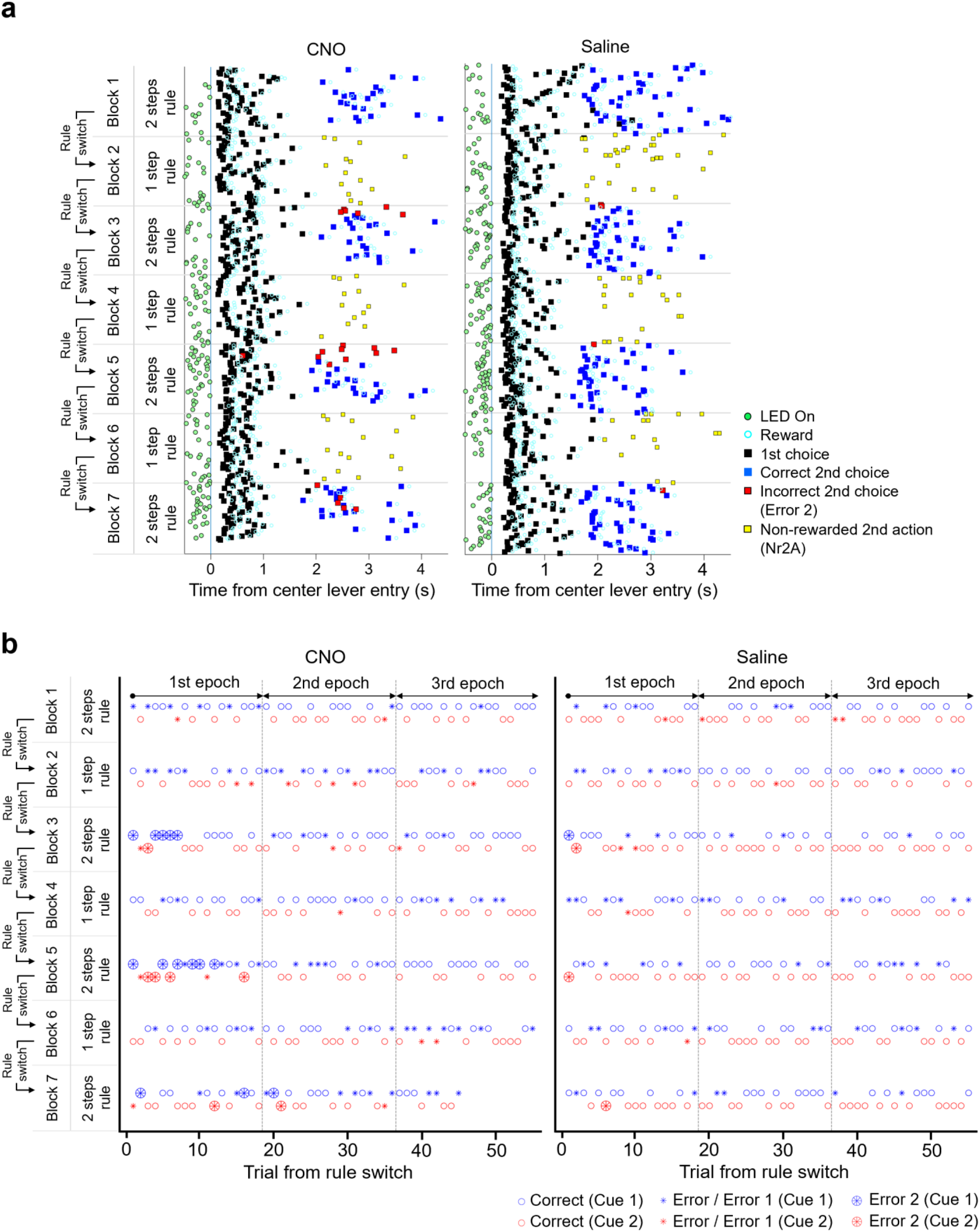
Task performance charts of representative sessions with chemogenetic silencing of ACC (see Fig. 2 main text and Extended Data Fig. 4). **a,** Events in individual trials in two representative sessions (same sessions as presented in Fig. 2b,c) with an *i.p.* injection of clozapine-N-oxide (CNO) solution (left, 10 mg/kg) or of a saline solution (right). Task events were sorted by the timing of animal’s pushing the center lever for initiating a trial. Green filled circle, LED onset. Black filled square, 1st choice response. Blue filled square, a correct 2nd choice response. Cyan circle, a water reward. Red filled square, incorrectly pushing the center lever before poking the opposite side port (“Error 2”). Yellow filled square, an entry to the opposite side port after making a correct 1st choice in 1 step rule condition for which no reward was delivered and instead a buzzer sound was presented (“non-rewarded 2nd action” or Nr2A). Trials were plotted from top to bottom (i.e., the first trial in a session was plotted at the top row). In every 55 trials, task rules were switched between 1 step and 2 steps rule conditions. **b,** Task performance chart of the same two representative sessions as in **a**. Small circle, correct trial. Asterisk, incorrect 1st choice (Error 1). Large circle filler with an asterisk, incorrect 2and choice trial (Error 2). Blue and red represent trials with two distinct tone cue stimuli. Gray dotted lines distinguish three epochs in each block (1-18th, 19-36th and 37-55th trials for 1st, 2nd and 3rd epoch, respectively).

**Extended Data Fig. 4.**
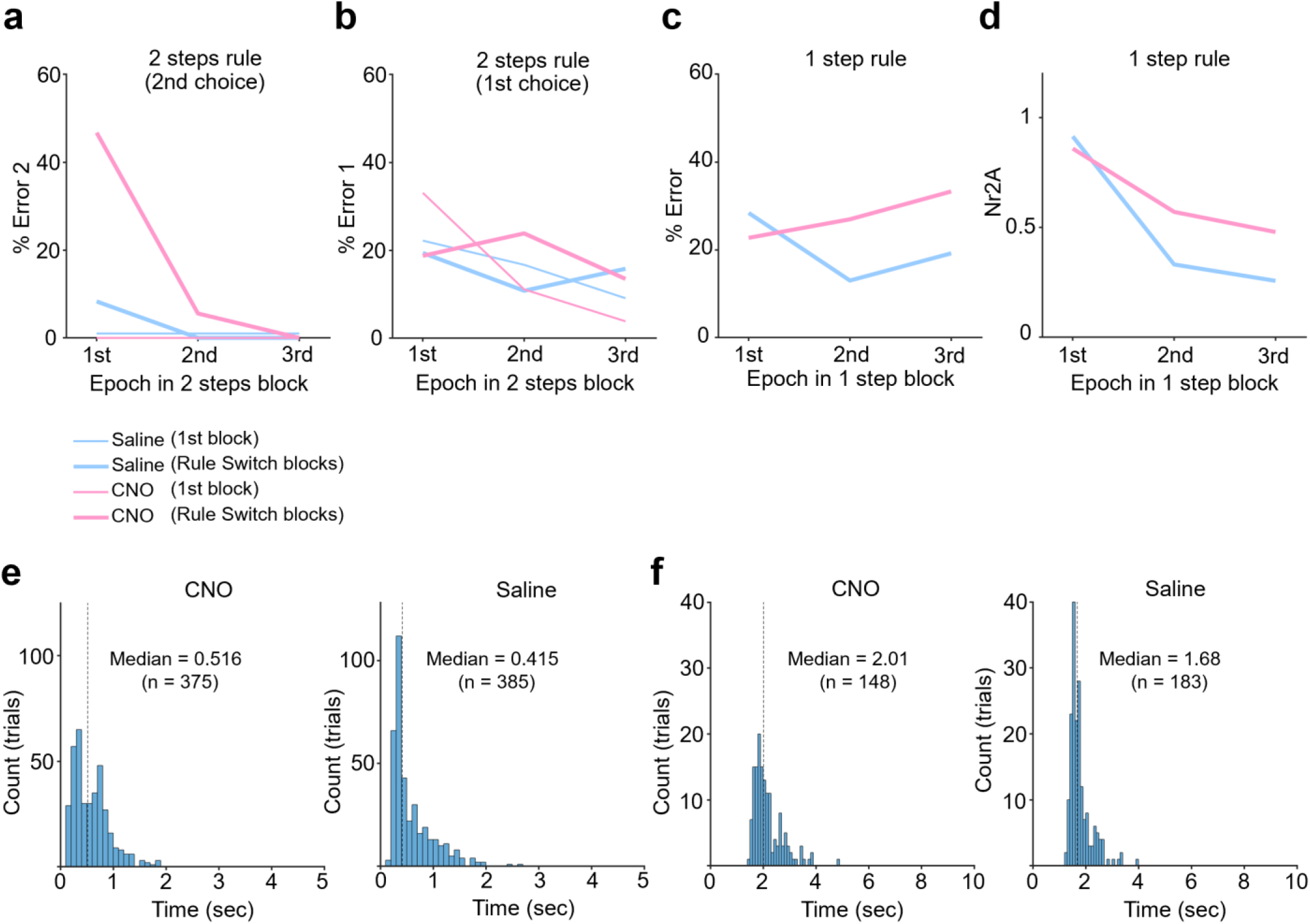
Behavioral effects of chemogenetic silencing of ACC in representative sessions (see Fig. 2 main text and Extended Data Fig. 3). **a,** 2nd choice performance in 2 steps rule condition (%Error 2) was plotted for two representative sessions (same sessions as presented in Fig. 2b,c and Extended Data Fig. 3) with an *i.p.* injection of CNO solution (10 mg/kg, pink) or of a saline solution (blue). Trials in 1st block (thin red and blue lines) and Rule Switch blocks (thick red and blue lines) were grouped into three epochs (“1st”, “2nd” and “3rd”) according to the trial no. in the corresponding blocks. 1st, 2nd and 3rd epochs correspond to 1-18th, 19-36th and 37-55th trials, respectively. % error rate for two tone cue conditions were averaged. See Fig. 2b for performance within the 1st epoch (the first 18 trials). **b**, Same format as in **a**, but average 1st choice performance (% Error 1) in 2 steps rule condition was plotted. **c**, Same format as in **a** and **b**, but average choice performance (% Error) in 1 step rule condition was plotted. **d**, Average number of non-rewarded 2nd actions (Nr2A) per trial in 1 step rule block was plotted for each of three epochs. See Fig. 2c for performance within the 1st epoch (the first 18 trials). **e**, Histograms of response time for the 1st choice. All trials from 1 step and 2 steps rule conditions were combined. **f,** Same format as in **e**, but for 2nd choices of correct trials in 2 steps rule condition.

**Extended Data Fig. 5.**
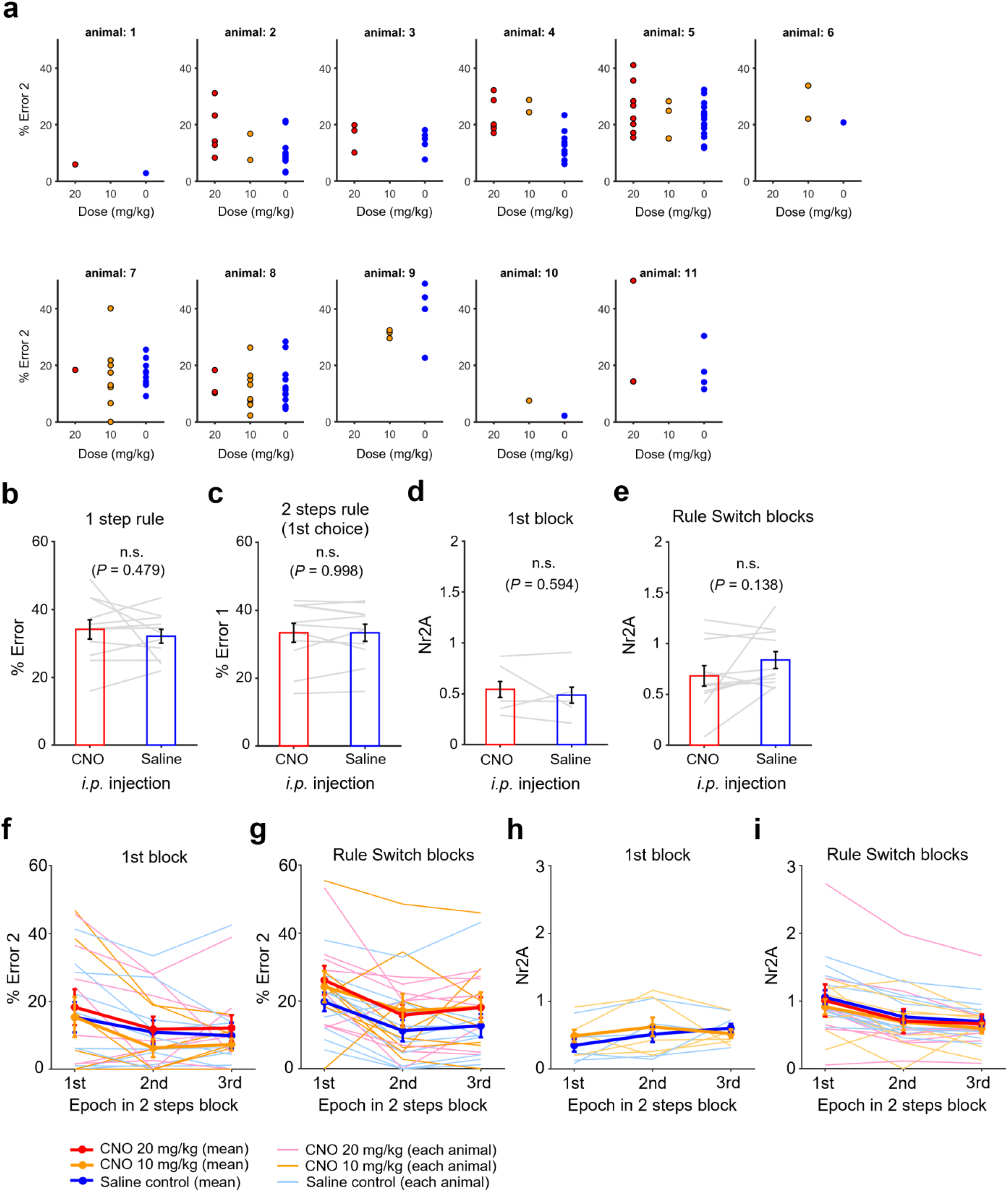
Group results of task performance with chemogenetic silencing of ACC neurons (see Fig. 2 main text). **a,** 2nd choice performance (%Error 2) in 2 steps condition for individual animals. Eight rats were tested in a CNO dose of 20 mg/kg (5 out of 8 rats were also tested in a CNO dose of 10 mg/kg). Additionally, another three rats were tested in CNO dose of 10 mg/kg. **b,** Group result of choice performance in 1 step rule condition (%Error) with an *i.p.* injection of saline or CNO solutions. For CNO data, sessions with doses of 10 mg/kg and 20 mg/kg were combined. Performance for two tone cues were averaged. Paired *t*-test, n = 11 rats. Error bars, s.e.m. **c**, Same as in **b**, but 1st choice performance (%Error 1) in 2 steps rule condition was plotted. **d**, Group result of average number of Nr2A per trial in the 1st block of 1 step rule condition. For CNO data, sessions with doses of 10 mg/kg and 20 mg/kg were combined. Performance for two tone cues were averaged. Paired *t*-test, n = 11 rats. Error bars, s.e.m. **e,** Same format as in **d**, but Nr2A was calculated for trials in Rule Switch blocks. **f**, 2nd choice performance (%Error 2) was plotted separately for three epochs (“1st”, “2nd” and “3rd”) in 1st block of 2 steps rule condition. Thick red, orange and blue lines represent across-animal averages of %Error 2 in 20 mg/kg dose of CNO, 10 mg/kg dose of CNO and saline conditions, respectively (n=8, 8 and 11 rats, respectively). Thin lines represent individual animals. 1st, 2nd and 3^rd^ epochs correspond to 1-18th, 19-36th and 37-55th trials. **g**, Same format as in **e**, but for Rule Switch blocks. **h**, Average number of Nr2A per trial was plotted separately for three epochs in 1st block of 1 step rule condition. Thick orange and blue lines represent across-animal averages of CNO and saline conditions, respectively. Thin lines represent individual animals (n=5 rats). 1st, 2nd and 3^rd^ epochs correspond to 1-18th, 19-36th and 37-55th trials. **i**, Same format as in **h**, but for Rule Switch blocks. Thick red, orange and blue lines represent across-animal averages of 20 mg/kg dose of CNO, 10 mg/kg dose of CNO and saline conditions, respectively (n=8, 8 and 11 rats, respectively). Thin lines represent individual animals.

**Extended Data Fig. 6.**
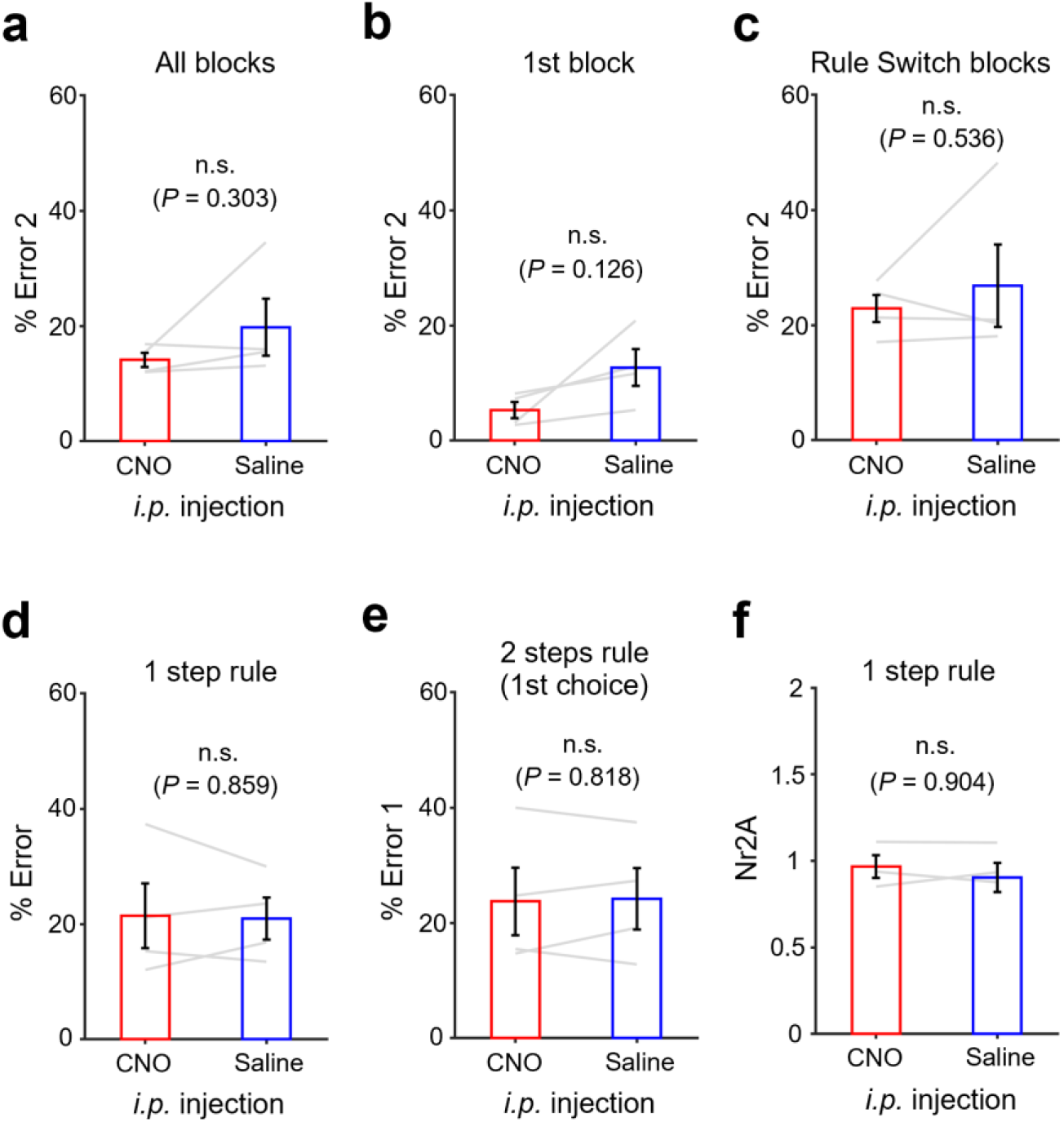
Administration of CNO showed no effect in choice performance in rats injected with control virus in ACC (see Fig. 2 main text). **a,** Group result of 2nd choice performance in 2 steps rule condition (%Error 2) with an *i.p.* injection of saline or CNO solutions in rats injected with AAV5-CamKIIa-mCherry virus in ACC. % error rate for two tone cue conditions were averaged. **b,** % Error 2 for trials in the 1st block (i.e., non-Rule Switch block) of 2 steps rule condition was plotted. **c,** Same as in **b**, but % Error 2 in Rule Switch blocks was plotted. **d,** Choice performance (%Error) in 1 Step rule condition was plotted (trials in 1st block and Rule Switch blocks were merged)**. e,** 1st choice performance (%Error 1) in 2 Steps condition (trials in 1st block and Rule Switch blocks were merged). **f,** Group result of average number of Nr2A per trial in 1 step rule condition. Paired *t*-test, n = 4 rats (**a**-**e)**, n = 3 rats (**f)**. Error bars, s.e.m.

**Extended Data Fig. 7.**
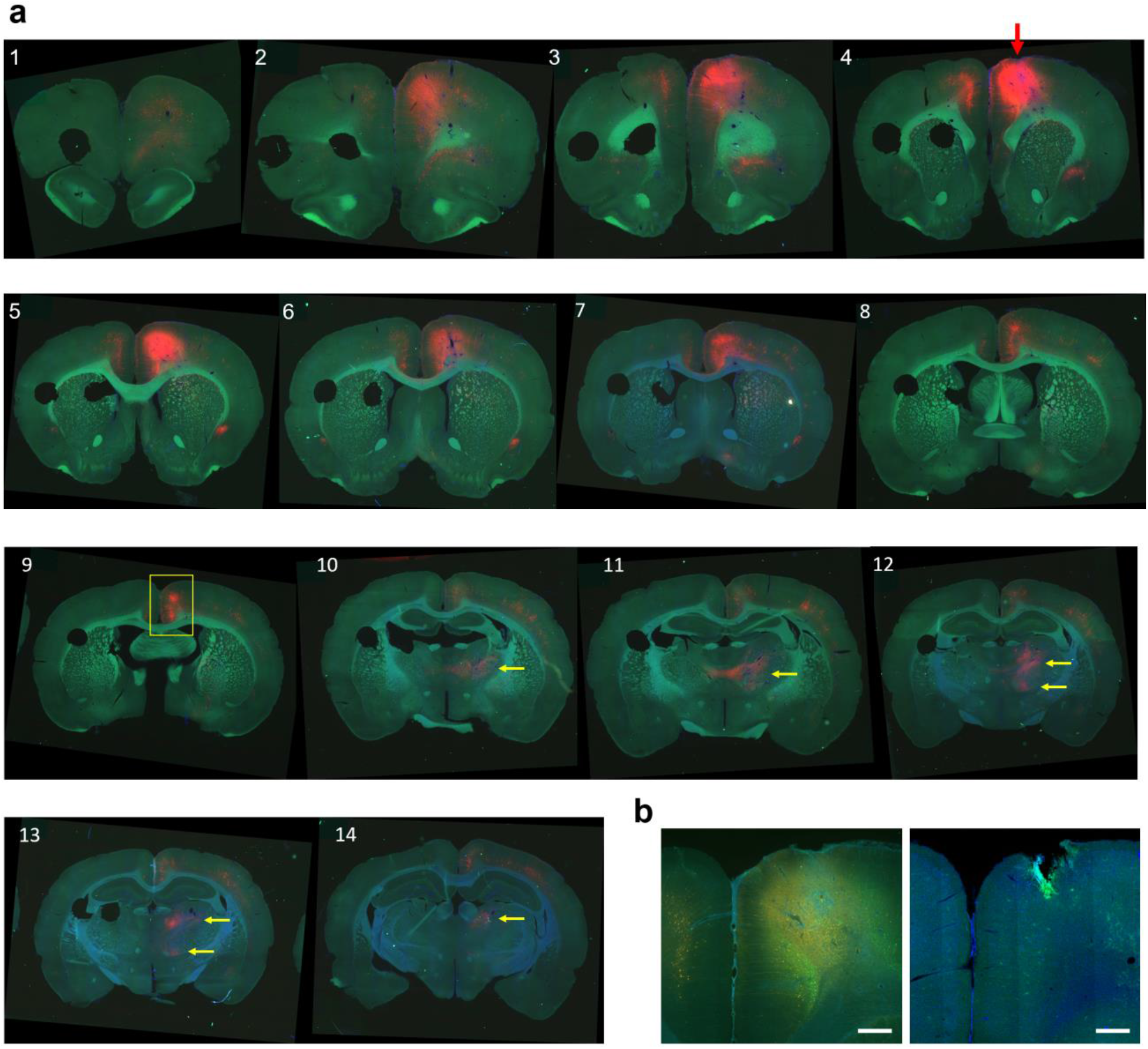
Cingulate and thalamic projections to secondary motor cortex (see Fig. 3 main text). **a,** Coronal section series of a rat’s brain. A cocktail solution (1 μl) of helper virus (AAV1- synP-FLEX-sTpEpB) and cre virus (pENN.AAV.CaMKII.0.4.Cre.SV40) was injected at secondary motor cortex (M2) followed by injection of genetically modified rabies virus (RV4-mChery (EnvA), 1 μl) at the same coordinate (pointed by a red arrow in panel no. 4). Neurons in anterior cingulate cortex (24a’/b’) were retrogradely infected with rabies virus expressing mCherry (indicated by a yellow square in panel 9, being also presented in Fig. 3a). Neurons in thalamic nuclei were also retrogradely labelled (pointed by yellow arrows in panels 10-14). Circular holes on the left side of sections were made before sectioning for the purpose of identifying the hemisphere contralateral to virus injections. **b**, Left, magnified view of the virus injection site in M2 (pointed by red arrow in **a**, panel no. 4) (this is an identical image as presented in Fig. 3b). Neurons that were infected by helper virus expressed GFP. Right, neither GFP nor mCherry expression was observed near virus injection sites in another wildtype rat in which helper virus was injected without AAV-CaMKIIa-cre virus. Scale bar, 500 μm.

**Extended Data Fig. 8.**
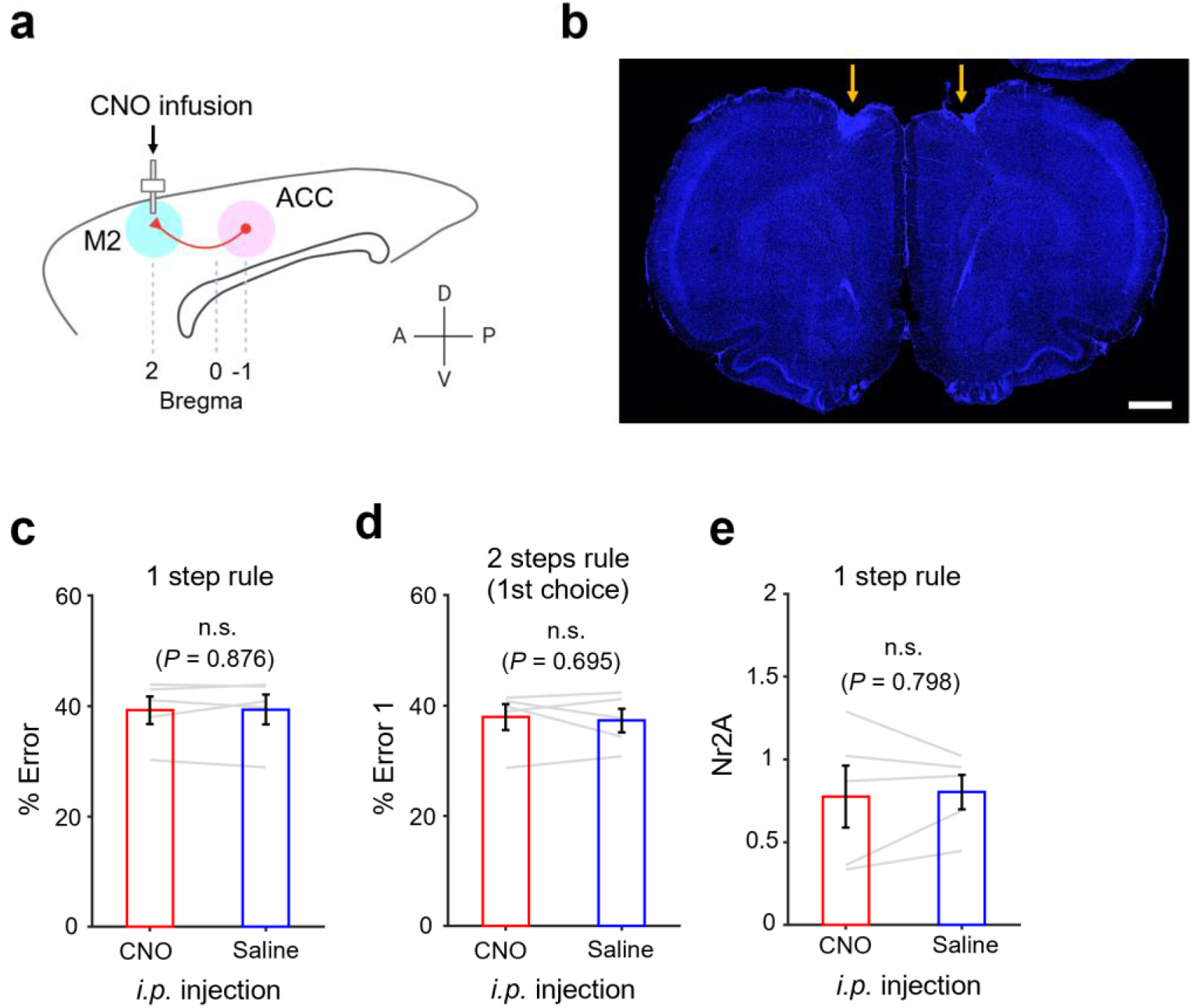
Chemogenetic silencing of ACC neuronal terminals in M2 disrupted the animals’ sequential choice performance after rule switches (see Fig. 3 main text). **a,** Rats were injected with AAV5-CaMKIIa-hM4Di-mCherry virus in ACC and were implanted with bilateral cannulae in M2. Animals’ task performance was tested with a local infusion of either saline or CNO solution (1 μg/μl) in M2. **b,** A coronal section showing tracks (orange arrows) of bilateral cannulae implant in M2. Scale bar, 1 mm. **c**, Group result of choice performance (% Error) in 1 step rule condition with local injection of saline or CNO solution in M2 (paired *t*-test, n = 5 rats). **d**, Group result of 1st choice performance (%Error 1) in 2 steps rule condition. **e,** Group result of average number of Nr2A per trial in 1 step rule condition. Paired *t*-test, n = 5 rats. Error bars, s.e.m.

**Extended Data Fig. 9.**
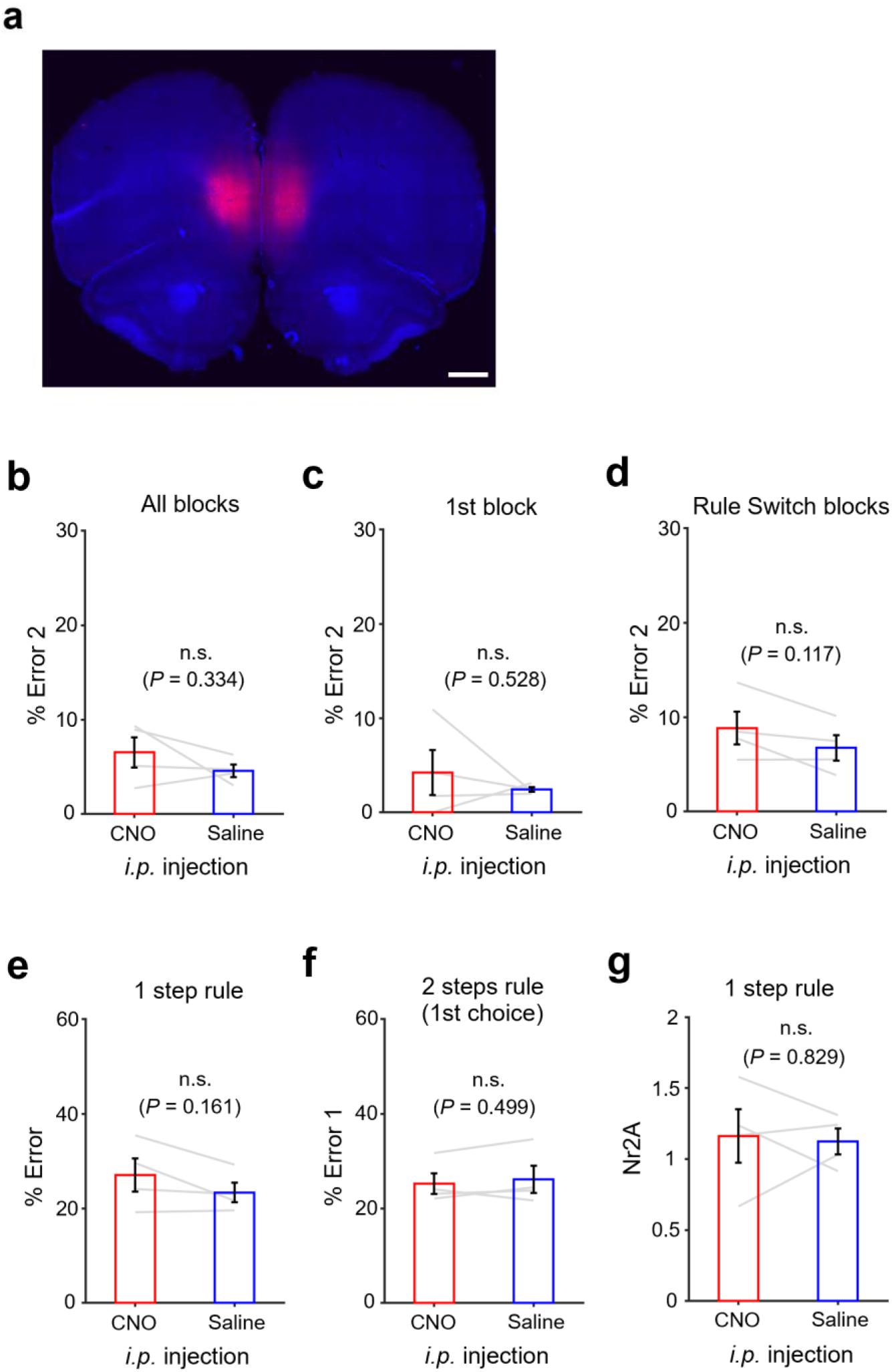
Chemogenetic suppression of prelimbic/infralimbic cortex showed no effect in choice performance. **a,** AAV5-CaMKIIa-hM4Di-mCherry virus (same virus as injected in ACC) was bilaterally injected in prelimbic/infralimbic cortex (1000 nl for each hemisphere). Scale bar, 1 mm. **b,** Group result of 2nd choice performance in 2 steps rule condition (%Error 2). % error rate for two tone cue conditions were averaged. **c,** %Error 2 for trials in the 1st block (i.e., non-Rule Switch block) of 2 steps rule condition was plotted. **d,** Same as in **c**, but %Error 2 in Rule Switch blocks. **e,** Group result of choice performance (%Error) in 1 step rule condition (trials in 1st block and Rule Switch blocks were merged)**. f,** Group result of 1st choice performance (%Error 1) in 2 steps rule condition (trials in 1^st^ block and Rule Switch blocks were merged). **g,** Group result of average number of Nr2A per trial in 1 step rule condition. Paired *t*-test, n = 4 rats. Error bars, s.e.m.

**Extended Data Fig. 10.**
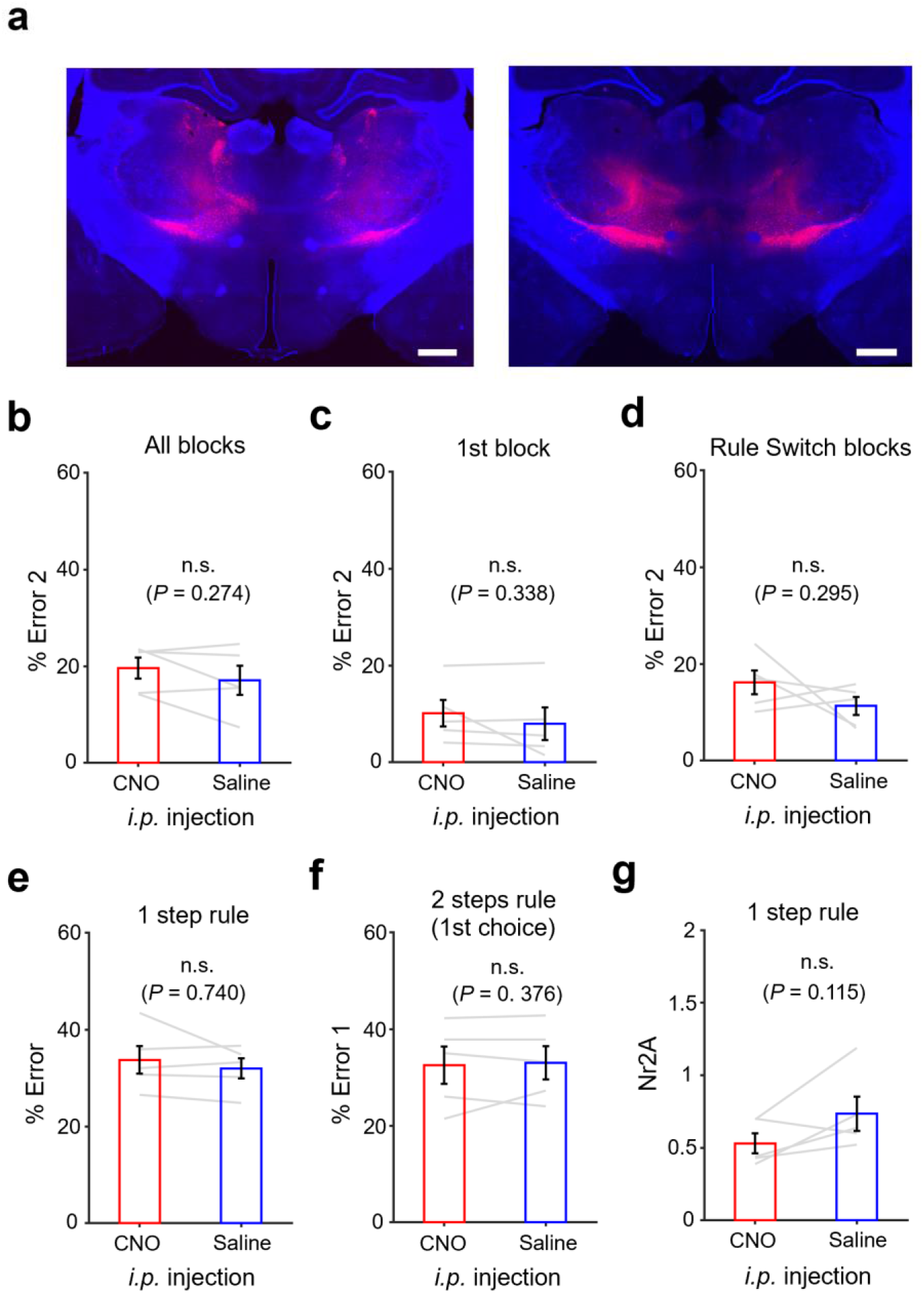
Chemogenetic suppression of ventral thalamus showed no effect in choice performance. **a,** AAV5-CaMKIIa-hM4Di-mCherry virus (same virus as injected in ACC) was bilaterally injected in ventral thalamic nuclei (1000 nl for each hemisphere). Scale bar, 1 mm. **b,** Group result of 2nd choice performance in 2 steps rule condition (%Error 2) with an *i.p.* injection of saline or CNO solutions. % error rate for two tone cue conditions were averaged. **c,** %Error 2 for trials in the 1st block (i.e., non-Rule Switch block) of 2 steps rule condition. **d,** Same as in **c**, but %Error 2 in Rule Switch blocks was plotted. **e,** Group result of choice performance (%Error) in 1 step rule condition (trials in 1st block and Rule Switch blocks were merged)**. f,** Group result of 1st choice performance (%Error 1) in 2 steps rule condition (trials in 1st block and Rule Switch blocks were merged). **g,** Group result of average number of Nr2A per trial in 1 step rule condition. Paired *t*-test, n = 5 rats. Error bars, s.e.m.

**Extended Data Fig. 11.**
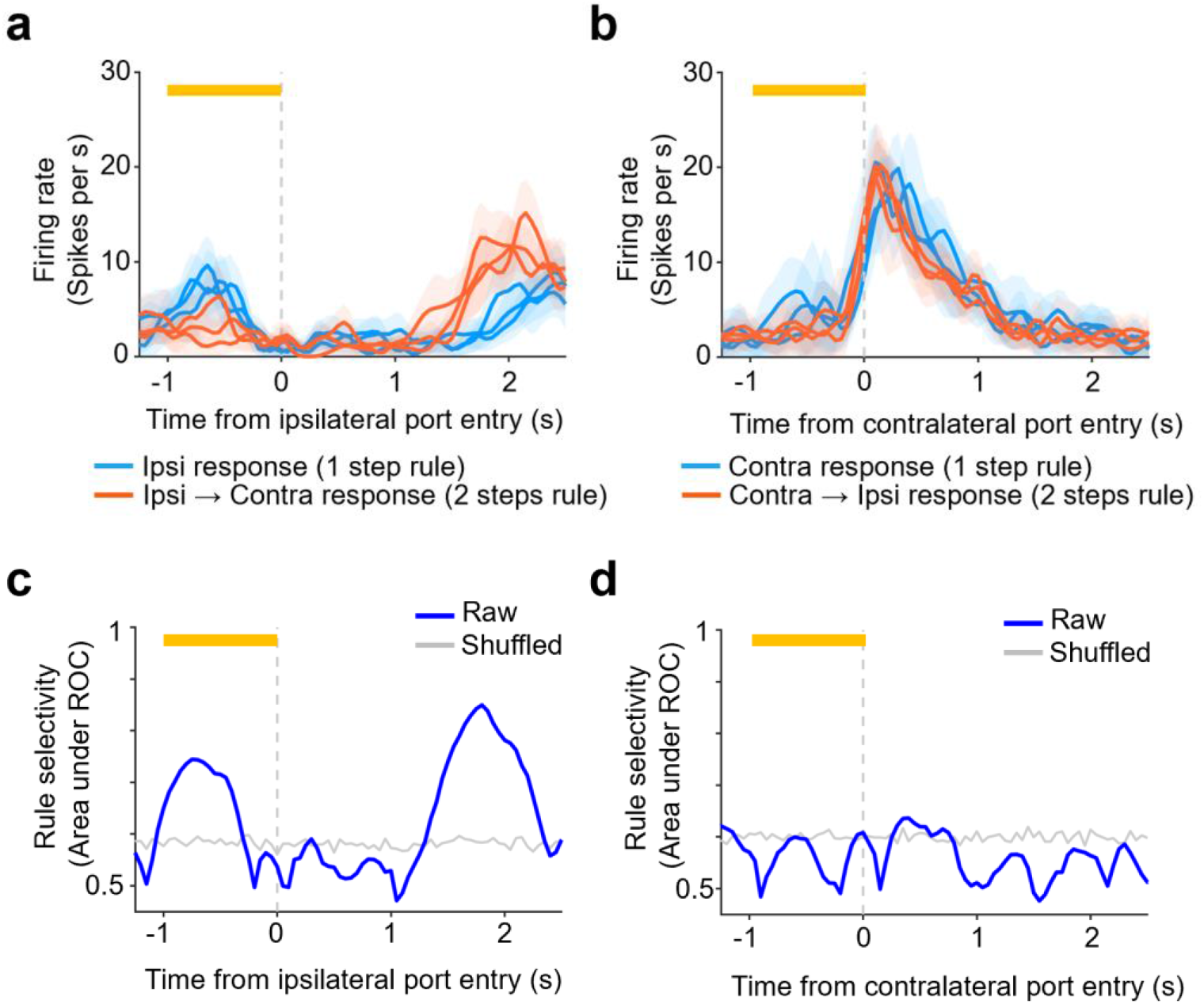
Rule selective activity of example M2 neuron preferring 1 step rule condition during pre-choice period (see Fig. 5 main text). **a,** Peri-event time histogram (PETH) of a representative single-unit showing rule selective responses preferring 1 step rule condition. PETH was calculated using trials in which the rat made a correct choice of ipsilateral side port, i.e., the side port that was located on the ipsilateral side of neural activity measurements. In 1 step rule block, the rat made a choice to the ipsilateral side port (blue line) while, in 2 steps rule block, the rat made a 1st choice to the ipsilateral side port and then made a 2nd choice to the contralateral side port (red line). Neural activity obtained in trials of Rule Switch blocks were plotted (note that neural activity in 1st block of the session was not included). Each line represents trial-averaged firing rates in a block (three blue lines and three red lines represent three Rule Switch blocks in 1 step rule condition and three Rule Switch blocks in 2 steps rule condition, respectively). Orange bar at the top, a 1 sec period immediately before rat’s entry to the ipsilateral side port (i.e., pre-choice period). Shaded bands, 95% confidence intervals. **b,** Same as in a, but for trials in which the animal made its correct choice response (1 step rule block) or correct 1st choice response (2 steps rule block) to contralateral side of neural measurements. **c,** Time course of rule selectivity of the single-unit presented in **a** and **b** was plotted for trials in ipsilateral condition. Gray line represents a 95% percentile level estimated by the shuffled data in which the area under ROC curve was calculated with rule labels for trials (i.e., 1 step or 2 steps rule conditions) being randomly shuffled. Error bar, s.e.m. **d,** Same as in **c**, but for trials in contralateral condition.

